# Bayesian optimisation and graph-based rheology enable sequence-dependent modelling of DNA materials

**DOI:** 10.64898/2026.05.27.728076

**Authors:** Aaron Gadzekpo, Lennart Hilbert

## Abstract

Bridging molecular and emergent properties is essential for designing soft matter. Synthetic DNA materials are attractive in this context because their sequence design space supports a wide range of material properties. Targeted design of DNA materials is hindered by scale differences and manual exploration of vast design spaces. We address this challenge with a computational workflow that links sequence-level design to rheological material properties. Concretely, we use machine learning to parametrise scalable, DNA-sequence-aware simulations, which we then evaluate using graph-based rheology. In our example, we study materials composed of self-interacting, multivalent DNA nanostars assembled from single strands. Structure and flexibility of nanostars are quantified with nucleotide-level oxDNA simulations, enabling Bayesian optimisation of a more coarse-grained bead-spring model. The bead-spring model allows efficient simulation of network formation between nanostars, governed by hybridisation free energies, which are computed with oxDNA and NUPACK. Nanostar valency and network connectivity are translated into rheological material properties with a graph-based method that we extend to include hydrodynamic interactions, yielding good agreement with experimental reference data. We generalise our findings by analysing theoretical graph representations of DNA materials and show how machine learning can optimise sequence affinities to produce desired rheological responses. Our work illustrates how machine learning can bridge scales and automate coarse-graining to facilitate targeted design of DNA materials through sequence-property relationships.

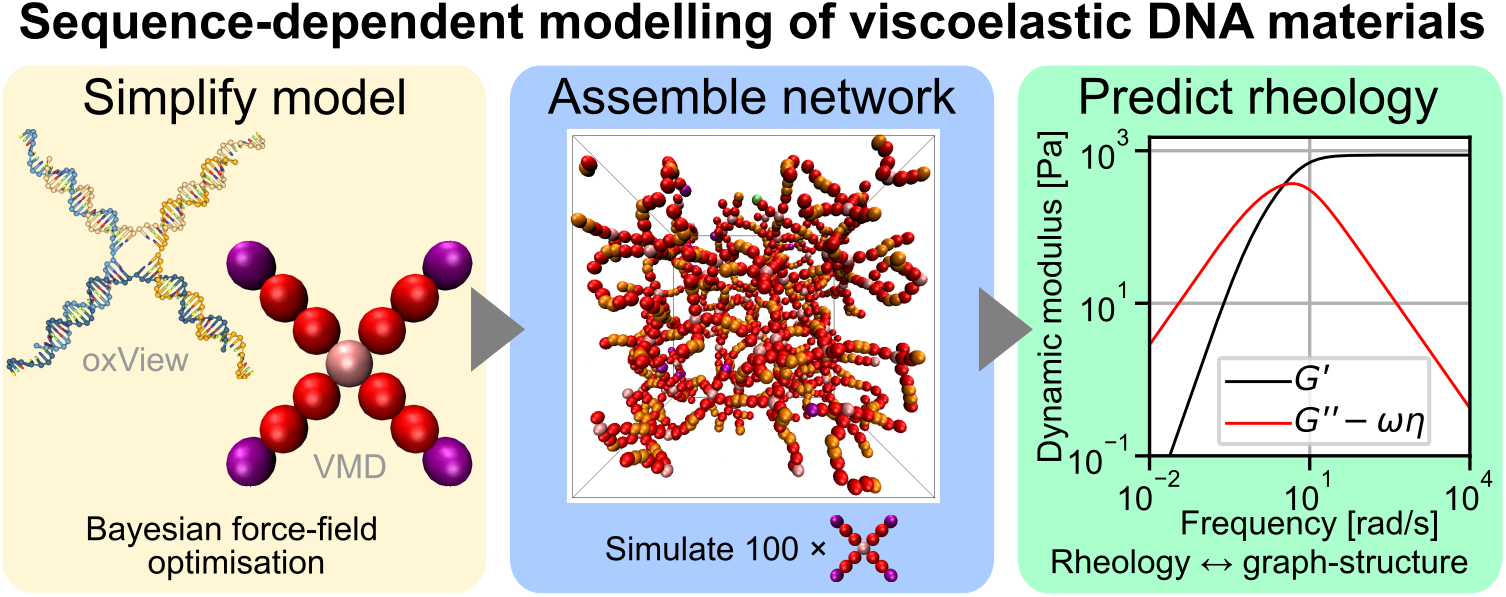

## 1 Introduction

DNA encodes genetic information through highly specific base pairing – a property that has been harnessed to construct synthetic 2D and 3D nanostructures. While static DNA origami [1, 2] has found diverse applications, [3, 4] dynamic and viscoelastic DNA assemblies, capable of responding to stimuli and being functionalised with other molecules, are emerging as powerful tools in biomimicry, biosensing, information processing, and biomedicine. [5– 16]

One emerging class of functional, dynamic DNA materials is based on networking of so-called DNA nanostars (also called DNA nanomotifs) into hydrogels and liquid-like condensates. DNA nanostars have multiple arms that can reversibly bind other nanostars via single stranded sticky ends or be linked with other molecules. [17–19] The design of these nanostars leads to a wide range of emergent behaviour such as (micro-)phase separation, reorganisation upon detection of other molecules, light or heat, as well as highly tuneable mechanical responses. [11, 20–24] This breadth of material properties opens new avenues for material design, given that the properties can be reliably engineered. Simulations and machine learning can accelerate this process by linking sequence-level design and emergent behaviour, guiding design choices while complementing and reducing the need for extensive experiments. [25]

To this end, simulations that resolve interactions at or below the nucleotide level are essential to investigate the microscopic origins of sequence-property relationships. However, simulations of these high-resolution models are limited in time and length scales. [26] Coarse-grained models, which group nucleotides into beads, offer a scalable alternative by sacrificing sequence specificity for computational efficiency. Yet, experimental evidence shows that mesoscopic properties, such as rheological behaviour, are highly sensitive to sequence-level changes, [11, 27] highlighting the need for multiscale approaches that connect nucleotide specific, microscopic properties with emergent, mesoscopic properties.

One promising route is to capture nanostar properties by sequence-specific optimisation of force fields for coarse-grained models. Efficient tuning of force field parameters can benefit from recent advances in machine learning, particularly Bayesian optimisation, capable of handling complex search spaces and minimal training data. [25, 28, 29] Here, we use Bayesian optimisation to refine a coarse-grained DNA nanostar model [25] that reproduces conformational statistics obtained from higher-resolution oxDNA [30–34] simulations. To complement structural analysis with sequence dependent binding affinities between DNA nanostars, we employ umbrella sampling in oxDNA. Further, we couple Bayesian optimisation with NUPACK [35–37] to predict DNA sequences and conditions for desired binding affinities. These thermodynamic insights integrate with the force field optimisation for a DNA sequence-aware modelling strategy. We implement nanostar models in ReaDDy, [38] a particle-based MD platform for reactive systems, enabling rapid exploration of nanostar network assembly within hours per design.

Network assemblies emerge through structure and affinity dependent nanostar-nanostar binding, producing mesoscopic rheological properties quantified by the storage (G’) and loss (G”) modulus. Considering that experimental data show moduli scaling from (near) Maxwellian well below the sticky end melting temperature [39, 40] to ∼ *ω*^1*/*2^ near the melting temperature [11, 41, 42], we wondered to what extend these networks can be described by a generalised Maxwell model. We therefore apply a method [43] that predicts the relaxation time spectrum of the generalised Maxwell model based on the material’s underlying graph structure, which we determine by assembling equilibrium networks with our optimised simulations. Additionally, we investigate the inclusion of hydrodynamic interaction between nanostars by employing an equilibrium approximation of the hydrodynamic interaction tensor, following Zimm’s extension of the Rouse model. Moduli are predicted up to pre-factors, which can be determined by extracting the experimental cross over point between storage and loss modulus. This limitation might be overcome with future work that links the pre-factors we set to unity to predictive values. Application of a graph-based approach complements existing computational methods by predicting the full frequency response directly from equilibrium structures, without requiring long trajectories of diffusing tracer particles or simulations of oscillatory shear over a wide frequency range.[27, 40, 44, 45] Our results align well with experimental data, outperforming the conventional Maxwell model across a broad frequency range and offering insight into the emergence of ∼ *ω*^1/2^ scaling near the gel point. To generalise our findings, we construct graph representations of other nanostar systems and explore their applicability and limitations by comparing to experimental data. The link between local molecular architecture, emergent network topology, and the resulting relaxation spectrum offers a microscopic explanation for experimentally observed deviations from the conventional Maxwell model. [46, 47] Our findings place DNA nanostar hydrogels within the wider context of colloidal and patchy-particle systems, in which fractal connectivity and valence polydispersity similarly control macroscopic rheology. [48, 49] To demonstrate potential inverse material design strategies, we combine simulations, graph-analysis, and Bayesian optimisation into a workflow that predicts sticky end sequences for nanostar networks producing prespecified G’ and G” curves. Our multiscale workflow connects sequence-level design to mesoscopic behaviour and integrates with any simulation framework with Python-accessible in- and output.

## 2 Methods

### 2.1 Nucleotide level simulations with oxDNA

For insights into sequence dependent properties of DNA nanostars, we used oxDNA, a nucleotide-level coarse-grained model that represents each nucleotide as a rigid body. [30–34] oxDNA captures base pairing, stacking, and backbone connectivity while implicitly modelling the solvent and was parameterised top-down to reproduce experimental data on DNA thermodynamics and mechanical behaviour. [31] The model has been widely applied in both biotechnology and fundamental DNA research. [50–55]

Using oxDNA, we carried out molecular-dynamics (MD) simulations of four-armed DNA nanostars assembled from 4 strands containing 49 nucleotides each. [17] Each strand ended with a 6-nucleotide sticky end at the 3’ end in the assembled nanostar that could bind to other sticky ends. An unpaired adenine at the start of the sticky end made it more flexible. [17] Nanostars were equilibrated and simulated in NVT ensemble for 5·10^7^ steps. To translate nanostar conformational flexibility into a simplified model, we grouped the nucleotides within each arm into coarse-grained beads. Six vectors aligned to these beads were used to generate angle distributions from the MD trajectories. The angles quantified flexibility within double- or single-stranded arm segments, and between arms. The angles reflected the degrees of freedom later parametrised in the bead-spring model. Simulation details and angle definitions can be found in the supplementary information (SI, Sections S1, S2).

Additionally, we performed MD simulations and umbrella sampling of the oxDNA model with a virtual-move Monte Carlo algorithm (VMMC) to obtain free energy profiles for two key processes: hairpin formation within sticky ends and hybridisation between arms. These thermodynamic profiles informed the binding behaviour in our coarse-grained model. To quantify the free energy associated with hybridisation events within and between sticky ends, we followed established protocols and defined two reaction coordinates for umbrella sampling: distance between nucleotides and presence of hydrogen bonds. Based on the frequency with which states defined by these coordinates were sampled with VMMC, weights were iteratively assigned to increase the frequency of sampling rare states, such as the binding of only one pair of nucleotides in the hairpin structure. [56] Free energy curves averaged six repeats of 10^7^ VMMC steps following equilibration. A statistical correction for finite size effects was used to derive bulk statistics from simulations of two sticky ends. [57]

### 2.2 Bead-spring model in ReaDDy

To simulate networks of 100 DNA nanostars, we used a bead-spring model with implicit solvent implemented in ReaDDy. [38] Each nanostar was represented by 13 beads, each bead approximating roughly 8 base pairs. Harmonic potentials defined bonds and angles between doublets and triplets of beads respectively. Salt dependent electrostatic repulsion between beads was modelled using a Debye-Hückel potential with a finite cut-off.

To capture sequence-, temperature-, and salt-dependent flexibility, [58–60] we treated the spring constants within arms and between bound arms as well as spring constants and equilibrium angles between adjacent or opposite arms as free parameters. This choice aimed to make the force field tunable for various experimental conditions.

The terminal bead on each arm represented a sticky end capable of hybridising with others via a reversible fusion reaction. In other words, we approximated the formation of hydrogen bonds between the sticky ends by stochastic addition and removal of a coarse-grained bond. This bond was modelled via a harmonic bond potential and an optimised angle potential. A separate conversion reaction toggled sticky ends between hairpin and open states. Bonds could be formed when to sticky ends were not in a hairpin configuration and close together in space. To capture the dependence of the bonded fraction of sticky ends on their DNA sequence and external parameters, we selected rates for hairpin formation based on hybridisation probabilities sampled with oxDNA. Furthermore, we tested a range of binding/unbinding rates to explore the influence of connectivity on mechanical properties in the material.

This approach allowed to select equilibria between binding and unbinding approximating experiments as well as unexplored ranges. Absolute rates were chosen to allow efficient simulation of gradual network assembly with fast hairpin formation and slower binding and unbinding between sticky ends. MD runs using Brownian dynamics were carried out for 5·5^5^ steps to sample angle distributions and for 1.5 − 6 · 10^7^ steps to assemble networks of 100 nanostars. Simulations started with an equilibration phase of 2 · 10^6^ steps without bond formation to relax the structure and remove unphysical initial configurations. Fractions of bound sticky ends and dynamic moduli were averaged from five independent repeats and configurations from the final 3.75·10^6^ time steps. Further simulation and parametrisation description can be found in the SI (Section S2).

### 2.3 Parameter tuning with Bayesian optimisation

To automate parameter tuning of our bead-spring model of DNA nanostars based on oxDNA reference data, we developed code that interfaced the in- and output of the simulation with the BoTorch library for iterative Bayesian optimisation. [61, 62]

The input in our workflow was a list of tunable simulation parameters with their respective ranges of permissible values, as well as distributions of reference observables, such as angles from high-resolution oxDNA simulations or target rheological properties. Each round of optimisation ran simulations with promising parameter combinations and extracted a fitness score by comparing the observed target quantity to the reference. The unknown relationship between input parameters and fitness was modelled using a Gaussian process trained on samples augmented with each optimisation round. The Gaussian process predicted mean fitness with associated uncertainty, guiding an acquisition function to propose new parameters likely to enhance fitness. This active learning strategy can balance exploitation (focus on high predicted mean fitness) and exploration (focus on regions of high fitness uncertainty) of the parameter space. [63]

As a fitness function for force field optimisation, we used an expression that increased in value for lower deviations between simulated and target angle distributions:

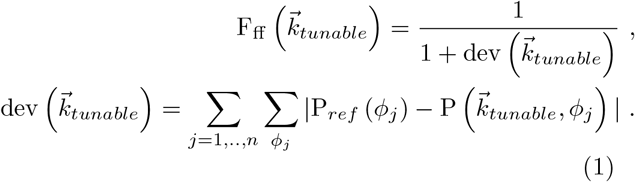

Here, 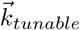 is a vector containing all tunable parameters that we wish to optimise to approximate the target angle distribution P_*ref*_ (*ϕ*_*j*_). The inner sum runs over the observed probabilities P (*ϕ*_*j*_) of angle *ϕ*_*j*_. When optimising parameters affecting different angle histograms, a sum over the histogram differences from each of the *n* angles was included.

For prediction of sticky end affinities based on target G’ and G” curves, we used a similar expression, quantifying the mean squared difference between observed and reference curves:

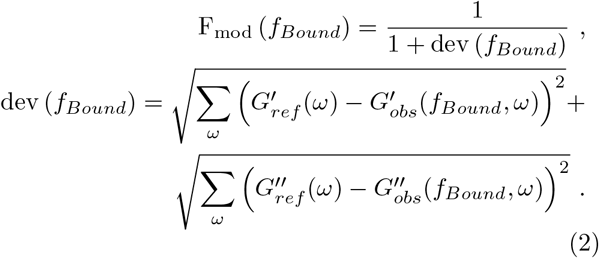

Here, *f*_*Bound*_ is the fraction of bound sticky ends in the material, which depends on sequence and conditions, captured in the simulations by (un-)binding rates.

These fitness functions produced values from 0 to 1, with the value 1 corresponding to the best fitness. BoTorch internally normalised the input parameters and standardised the fitness values, limiting the influence of the concrete choice of fitness function and absolute input values.

To improve statistics and accelerate optimisation, our implementation ran and evaluated five independent simulation instances in parallel. We also included the option to use separate Gaussian processes to model parameter-fitness relationships in parts of the simulated nanostars that do not interact strongly with each other. This strategy allowed, e.g., to optimise the force field in central parts of the nanostars simultaneously with the force field acting on nanostar sticky ends, by using separate Gaussian processes on the same set of trial simulations. This strategy aimed to reduce the number of optimisation steps and the dimension of the parameter search space, by factoring the search space into independent subspaces.

### 2.4 Sticky end optimisation with Bayesian optimisation

To find sticky end sequences and conditions that yield a target fraction of hybridisation, we interfaced NU-PACK’s test tube tool with Bayesian optimisation in BoTorch.

Sticky end sequences of N nucleotides length were represented as N-dimensional hypercubes, where the quartiles along each dimension encoded for one the four bases. Three additional dimensions encoded temperature and concentrations of mono- and bivalent salt ions. Sticky end length and concentrations were kept fixed. A test tube was set up for each design and NUPACK predicted what fraction of sticky ends would form a complex of size two. This hybridised fraction was compared to a target fraction *f*_*target*_ via the following fitness function:

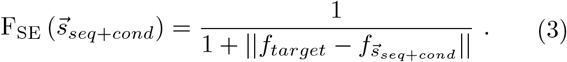

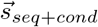 contains the sequence and conditions, producing a fraction 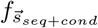 . Bayesian optimisation was then used as discussed for force field optimisation to predict a library of sticky end designs and conditions given a target fraction and permissible ranges for temperature and salt concentrations. Sequences can also be designed within NUPACK, [64, 65] here we opted for Bayesian optimisation for greater flexibility and to include tuning of salt and temperature conditions for more precise optimisation of binding equilibria.

### 2.5 Extracting rheological properties via the graph-Laplacian

To characterise the viscoelastic behaviour of DNA nanostar networks, we start out with the conventional Maxwell model, which describes materials with both elastic and viscous responses. This model corresponds to an elastic spring and a viscous damper in series, [47, 66] the storage modulus G’ and loss modulus G” are given by:

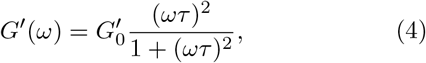

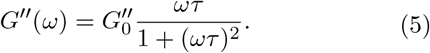

Here, *ω* is the angular frequency, and *τ* is the relaxation time.

Transitioning to polymer materials, the relaxation process can be interpreted in terms of the Rouse [67] or Zimm models [68] for polymer dynamics, where a spectrum of relaxation times emerges as sections of a polymer with different length relax with different time constants. The equation of motion for polymer bead *i* can be written as: [69, 70]

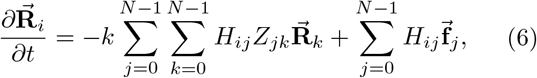

Where 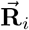 contains the positions of the polymer beads connected with spring constants *k, Z*_*jk*_ encodes the connectivity between polymer beads, the entries *H*_*ij*_ of the mobility matrix **Ĥ** account for hydrodynamic interactions, and 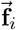 contains the stochastic force vectors. Replacing **Ĥ** with an identity matrix times a scalar factor 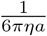 (the inverse friction coefficient 1*/ζ*) effectively results in the Rouse model, while replacement of **Ĥ** with its equilibrium average results in the pre-averaging approximation for polymer dynamics near equilibrium proposed by Zimm. [68, 69] Using the pre-averaging approximation, the elements *H*_*nm*_ are given by:

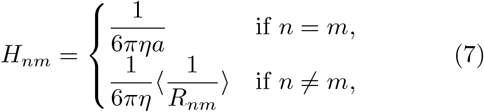

with an effective bead radius *a*, a medium viscosity *η* and the equilibrium average of the inverse distance 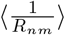 be-tween beads *n* and *m*. This approximation replaces the configuration-dependent hydrodynamic interaction with its equilibrium ensemble average, assuming that the fluid drag is determined by the mean polymer shape; this simplifies the dynamics into a linear system of independent relaxation modes. Since the *x, y, z* components are de-coupled, [71] the dynamics are described by N equations for a vector of modes 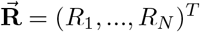 :

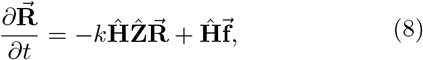

This equation may be solved by diagonalising the *N*×*N* matrices 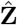or 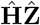 when hydrodynamics is considered. [68, 69] The eigenvalues obtained through diagonalisation are inversely proportional to the relaxation times, which leads to a generalised Maxwell model for storage and loss moduli: [43, 71, 72]

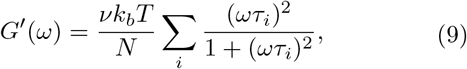

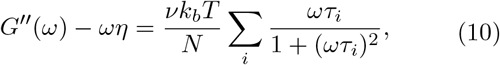

where (assuming dilute conditions) *v* is the number of beads per unit volume, *k*_*b*_*T* is the thermal energy an *N* the number of connected beads. G” now features a contribution from the solvent with viscosity *η*. The sum runs over all relaxation times

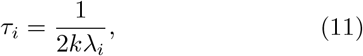

where *λ*_*i*_ are the eigenvalues of 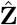 or 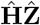 and a factor 2 accounts for the second moment of displacement involved in stress relaxation. [69] To analyse scaling predictions, all material prefactors were later set to unity.

The Rouse model was recently applied to protein condensates with intra- and intermolecular connectivity, [43, 73] motivating us to examine its applicability to hydrogels formed by DNA nanostars. In parallel, we explore predictions from a purely graph-theoretical perspective, modelling materials as networks of nodes with a maximum valency, where a tunable fraction of connections forms cross-links. We further incorporate hydrodynamic interactions in an approximate manner by extending the framework toward the Zimm model.

Figure 1A shows how scale-bridging and graph-based analysis for nanostar networks allowed to extract rheological properties. In networks of nanostars simulated with the optimised bead spring model, each motif represented a node and hybridised sticky ends added edges. The graph Laplacian 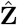of the example network is shown in figure 1B. Connections between nodes i and j result in an entry of -1 in the graph Laplacian matrix. The main diagonal contains the absolute number of connections terminating in each bead. The relaxation times extracted from this graph Laplacian are plotted in 1C and produced G’ and G” curves shown in 1D. There are three primary scaling regimes: the low frequency regime where G’∼*ω*^2^ and G” ∼*ω*, the intermediate regime where G’, G” ∼*ω*^*n*^ with 0 *< n <* 1 and the high frequency regime where G’ is constant and G” ∼*ω*^*−*1^. The exact scaling of the curves and the frequency ranges of the different regions are influenced by the connectivity in the polymer material. We treated all connections between nanostars as equivalent (no edge-weighting) and omitted nanostar-internal relaxation dynamics as well as contributions from topological linking [40] in the network. The graph Laplacian is time dependent as connections between sticky ends could stochastically split and fuse during the simulation. Hydrodynamic interactions can be approximated by multiplying the graph Laplacian with the mobility matrix defined in equation (7), where 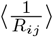 was extracted by averaging the inverse nanostar centre-centre distances over 20 frames from networks assembled and equilibrated with the bead-spring model. These networks, produced by extracting graph configurations from many time points and repeats, contributed to the final, averaged moduli.

**Figure 1.**
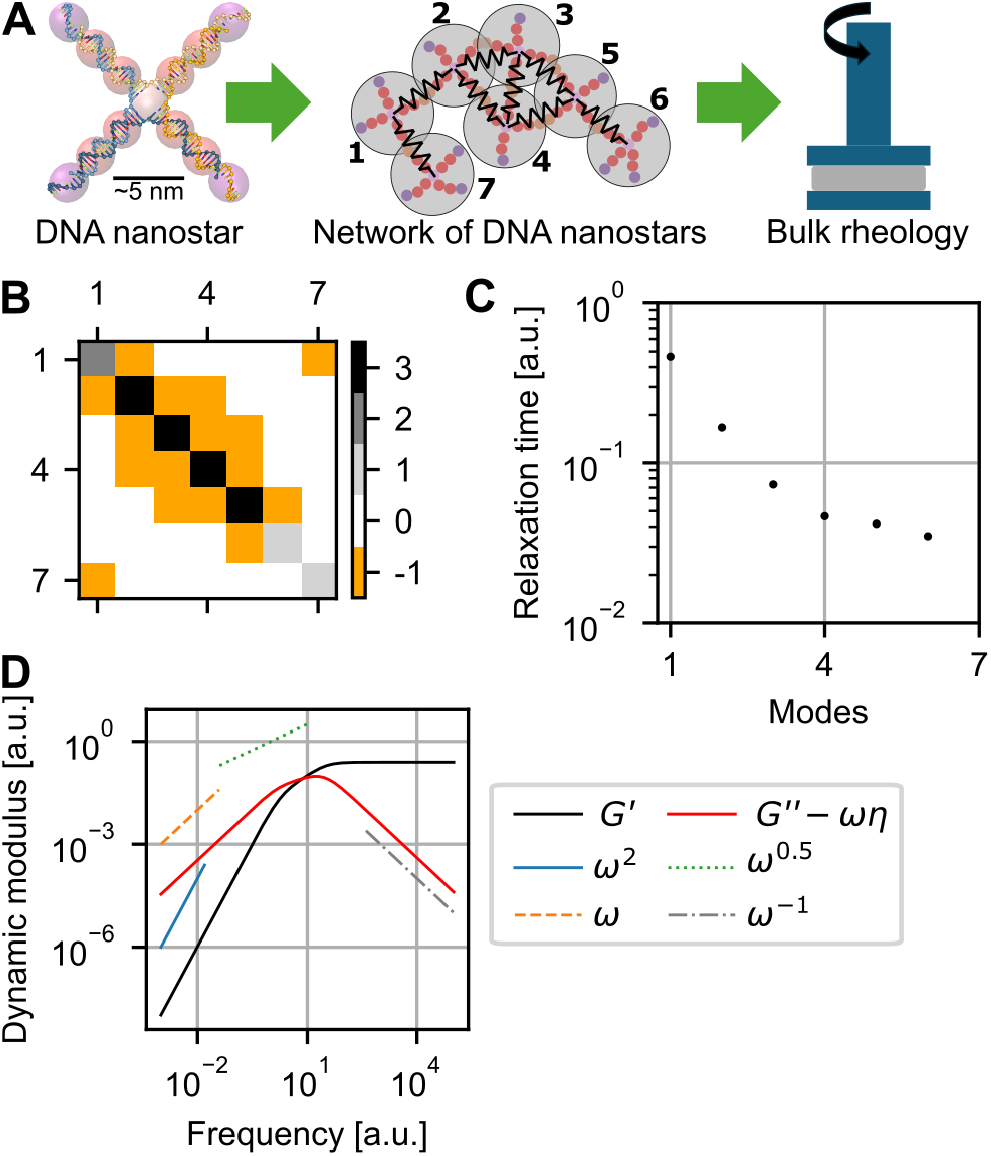
Method integration for sequence-to-rheology modelling. (A) DNA nanostars are the building blocks for viscoelastic materials analysed in this work. Machine learning bridged from nucleotide details to coarse-grained bead-spring models. Treating entire nanostars as nodes in a network allowed to connect simulated assemblies to rheology measurements. (B) Graph Laplacian of the nanostar network in A. The connections between nanostar nodes i and j were represented by an entry of -1. The main diagonal summed the connections terminating in each node. (C) The spectrum of relaxation times corresponding to the graph Laplacian in B. Relaxation times are inversely proportional to the eigenvalues of the graph Laplacian. (D) The relaxation times were used to calculate the storage modulus G’ and loss modulus G”. In the low frequency regime, G’*∼ ω*^2^ and G”*∼ ω*. In the intermediate regime, G’, G”*∼ ω*^*n*^, 0 *< n <* 1. In the high frequency regime, G’ becomes constant and G”*∼ ω*^*−*1^. The shape of both curves as well as the widths of the scaling regimes depends on the polymer connectivity.

## 3 Results

### 3.1 Quantifying nanostar flexibility

Starting at the nucleotide-level, we used oxDNA simulations to capture conformational statistics of single and paired DNA nanostars. In brief, we evaluated the overall shape of the nanostar and then quantified structure and flexibility by aligning vectors to groups of nucleotides. Distributions of angles between these vectors then served as optimisation targets for the sequence-informed bead-spring model.

In oxDNA simulations, the nanostar switched over time between three configurations resembling a flat X: one with an open central structure (configuration *γ*) and two with a cross of double-stranded DNA at the centre (configurations *α* or *β*) (Figure 2A). These three configurations could be differentiated by comparing combinations of angles between the four arms. Labelling the arms A, B, C and D, angle A&D refers to the angle between vectors aligned to arms A and D, and the same applies to the other combinations. When comparing the time evolution of the sum of angles A&D and B&C to the sum of angles A&B and C&D, one can differentiate between the three configurations (Figure 2A). If one sum was significantly larger than the other, the motif was in configurations *α* or *β* with crossing double-stranded DNA at the centre. If the two sums have similar values, the motif was in configuration *γ* with open internal structure. Under the tested conditions, the nanostars were almost exclusively in states *α* or *β*, with occasional switches between them (Figure 2A). Note that this strongly depended on salt concentration and temperature. Differentiating between the three configurations allowed to assign a “base angle” corresponding to the angle between adjacent arms separated by *≈* 90° and an “opposite arm angle” corresponding to arms whose sticky ends point in opposite direction (≈180° separation) (Figure 2B). This step was necessary as the bead-spring model represented an averaged structure that did not resolve occasional switching between the three states. Besides the two angles capturing angle correlations and fluctuations between arms, we also defined angles quantifying the flexibility within the double-stranded arm section (“Arm axis angle”), between two arms fused by sticky end hybridisation (“Bound angle”) and between the double-stranded arm and the sticky end, which is either open and single stranded (“Link angle”) or partly double-stranded when forming a hairpin (“HP link angle”). The resulting six angle distributions (Figure 2C) provided a reference for the flexibility of the bead spring model.

**Figure 2.**
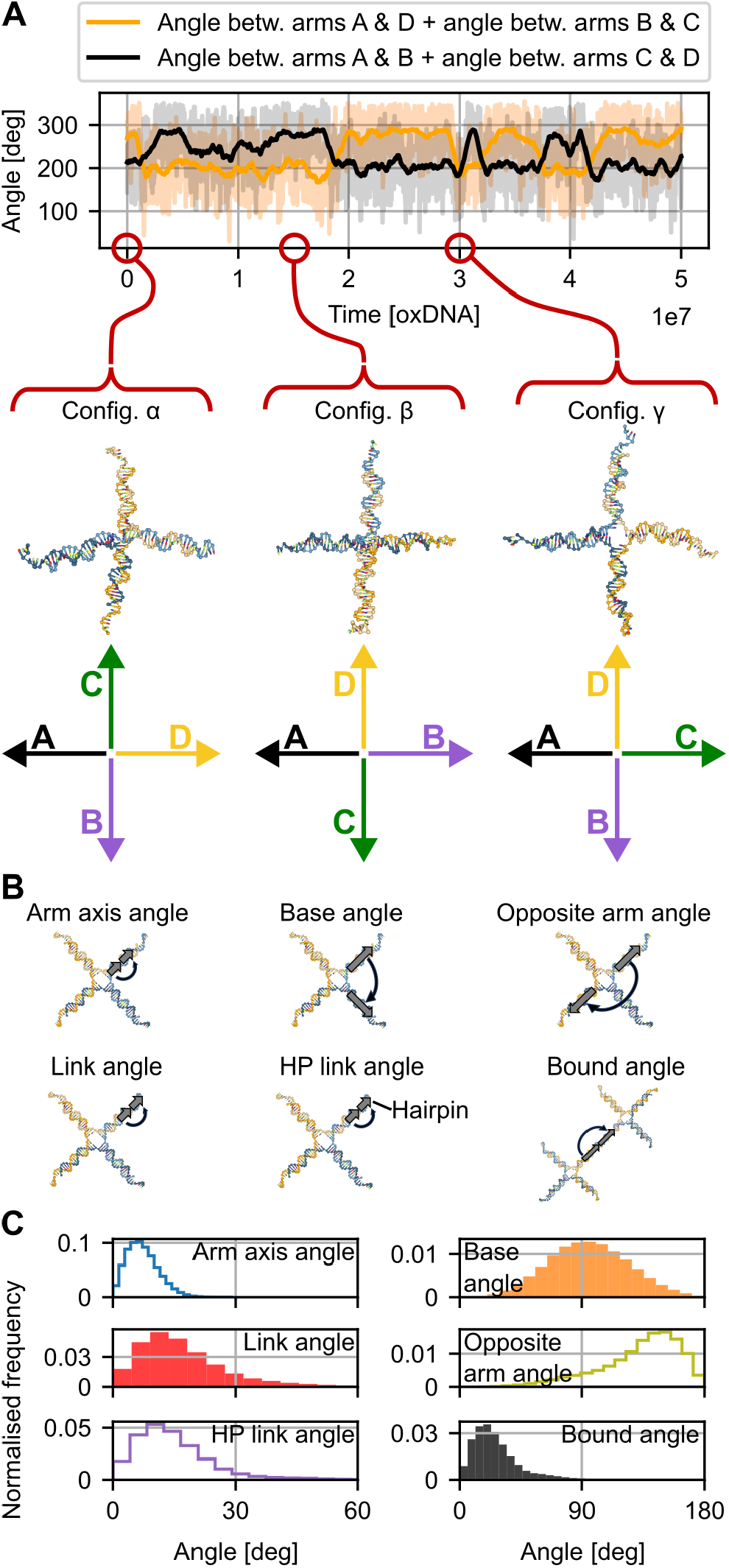
Capturing nanostar angles. (A) Assigning fixed labels A, B, C and D to the four nanostar arms allowed to define four angles between arms. Plotting the time evolution (moving average in darker shade) of two sums of angles revealed switching between configurations *α, β* and *γ*, which differ in the structure at the centre of the X shape. (B) Six angles within and between nanostar arms were defined by aligning vectors to group of nucleotides. (C) Resulting angle distributions obtained from MD simulation.

### 3.2 Bayesian optimisation of the bead-spring model

Using Bayesian optimisation, we tuned force field parameters of a bead-spring model of DNA nanostars to reproduce target angle distributions. We first validated the approach by using random parameter combinations and resulting angle distributions as optimisation targets. Following validation, we used target angle distributions from oxDNA to parametrise our model for subsequent use.

To reproduce angle distributions that could vary with nanostar design and physicochemical conditions, six angle potentials with eight free parameters were tuned. Angle potentials were defined between three adjacent beads along nanostar arms or across the nanostar centre (Figure 3A).

**Figure 3.**
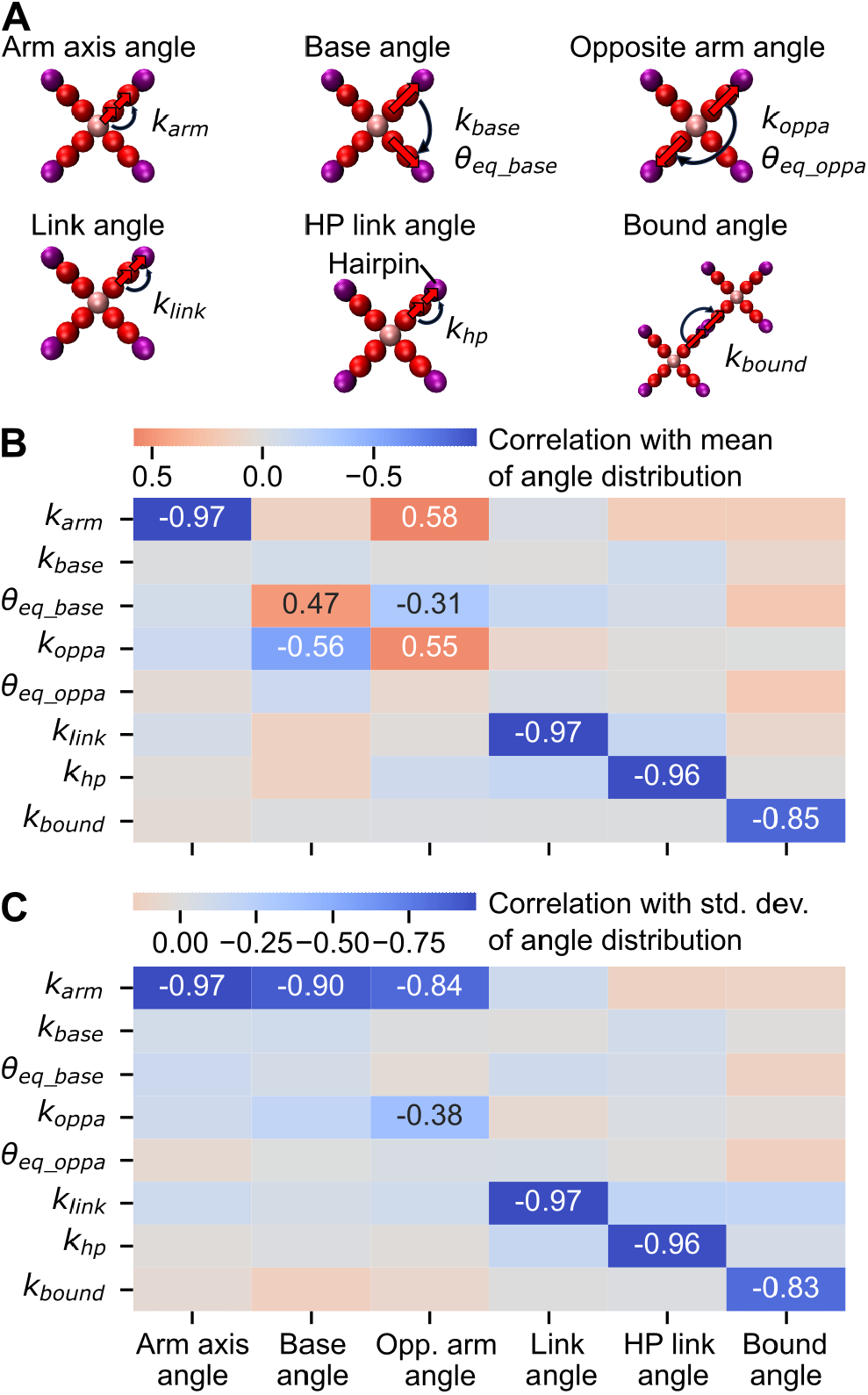
Correlations between parameters and angle distributions. (A) Eight parameters of angle potentials and six resulting angle distributions were evaluated in the bead-spring model. Spearman correlation between parameters and the mean (B) and standard deviation (C) of the resulting angle distributions. Angle distributions were averaged from five repeats of simulations with 100 random parameter sets.

To examine correlations between the eight free parameters and resulting angle distributions, we simulated 100 random parameter sets across a broad range of permissible values (SI, Table S1). Spearman correlations between parameters and the mean and standard deviation of angle distributions can be found in Figure 3B,C. The correlation patterns revealed that varying the spring constant within arms *k*_*arm*_ affected the angle along the arm (“arm axis angle”) as well as the angles between adjacent and opposite arms (“base angle” and “opp. arm angle”). The angles between adjacent and opposite arms were also influenced by the spring constant and equilibrium angle *k*_*oppa*_ and *θ*_*eq_base*_ but only weakly by *k*_*base*_ and *θ*_*eq_oppa*_. These couplings between parameters suggested that serial optimisation of individual parameters would be inefficient, motivating two joint strategies:

1. Split optimisation: optimise *k*_*arm*_ first, capturing the flexibility of double-stranded DNA, then jointly optimise *k*_*base*_, *k*_*oppa*_, *θ*_*eq_base*_, *θ*_*eq_oppa*_, capturing interactions between arms
2. Combined optimisation: optimise *k*_*arm*_, *k*_*base*_, *k*_*oppa*_, *θ*_*eq_base*_, *θ*_*eq oppa*_ simultaneously, leading to a larger search space.

Spring constants for open or hair-pinned sticky ends (*k*_*link*_, *k*_*hp*_) and bound nanostar arms (*k*_*bound*_) were optimised separately following split or combined optimisation, as they correlated with their corresponding angle distributions and only weakly with other angles. To reduce the number of optimisation simulations, *k*_*link*_ was optimised with a second Gaussian process using the trial simulations of the multi-parameter optimisation.

We evaluated the performance of both strategies on 13 random parameter sets in the bead-spring model. Split optimisation used 16 initial and 120 optimisation iterations for the 4 jointly optimised parameters. Combined optimisation used 32+120 iterations for the 5 jointly optimised parameters. Single parameter optimisation used 2 initial and 20-30 optimisation iterations. Initial simulations explored extreme cases by sampling the corners of the parameter space. The number of optimisation iterations typically lead to convergence and was kept constant for better comparability (SI, Figure S4). Performance was assessed via Wasserstein distance and Kolmogorov-Smirnov (KS) tests between optimised and target angle distributions (SI, Figure S5). Both strategies performed similarly, with split optimisation overall being more reliable. Optimised angle distributions matched their targets visually and according to KS tests in *≈* 80-100% of cases. Examples of (un-) successful optimisation and corresponding Wasserstein distances as well as overall performance evaluation can be found in the SI (Section S3). We then applied the method to oxDNA targets, using the same strategies and number of iterations used during validation. Combined optimisation performed notably worse than split optimisation on the base, opposite and bound angle, but almost identical for all other angles (SI, Figure S6). Split optimisation produced a close match between optimised and target angle distributions (Figure 4). Only the base and opposite angle distributions showed obvious deviations from their targets, likely due to the much-reduced complexity of the bead spring force field at the central nanostar structure. The split optimisation strategy converged on a solution that set *k*_*base*_ to the lowest permissible (effectively zero) value, indicating that this degree of freedom was not needed to match the oxDNA targets. This result may also reflect the fact that *k*_*base*_ only weakly correlated with any angle distribution (Figure 3B,C). Optimised spring constants ranged from stiff arm potentials (*k*_*arm*_ = 2.033 · 10^*−*19^ J*/*rad^2^) to more flexible bound arm potentials (*k*_*bound*_ = 1.572 · 10^*−*20^ J*/*rad^2^) and even more flexible opposite arm potentials (*k*_*oppa*_ = 5.836 10^*−*21^ J*/*rad^2^). These differences reflected the underlying structural context: fully double-stranded regions were stiffer than sections that were partially single stranded, or regions separated by the flexible, partially unhybridised centre section. A complete list of optimised parameters is provided in the SI (Table S1).

**Figure 4.**
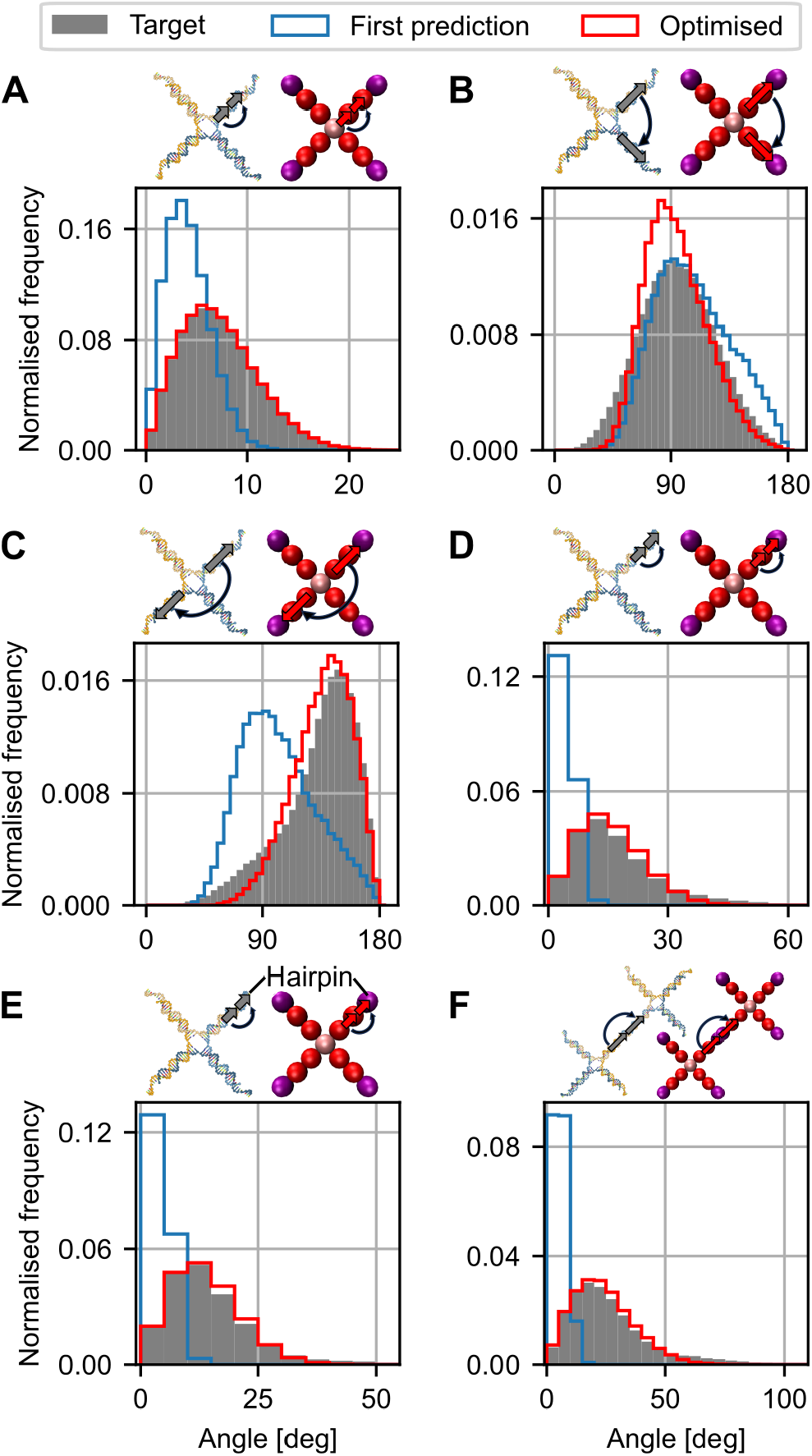
Optimised angle distributions. (A)-(F) First prediction after initial simulations and optimised angle distributions, compared to oxDNA targets. Angle distributions from bead-spring model averaged from five repeats.

We conclude that careful factorisation of the parameter space and informed choice of the optimisation order produced angle distributions closely approximating their targets, while compromising between distributions that are strongly coupled.

### 3.3 Hybridisation within and between nanostar arms

To complement the tuning of mechanical properties carried out previously, we quantified the thermodynamics of reversible hybridisation between nanostar sticky ends with oxDNA and NUPACK. In short, free energies and binding equilibria informed the fusion and fission reactions that modelled sticky end hybridisation in our bead-spring model.

The sticky ends had a palindromic sequence, allowing them to form hairpins. We compared umbrella sampling of hairpin formation in the seven nucleotides of the sticky end to MD simulations of the entire nanostar. Umbrella sampling produced free energy curves almost identical the MD simulation, when an additional six base pairs of arm segment were included (SI, Figure S3). Including an arm segment likely captured the reduction in entropy caused by constraining one side of the sticky end. The full free energy profile as a function of base pairs present in the hairpin can be seen in Figure 5A. At 20°C, our target temperature for comparison to experiments, the fully open configuration had the lowest free energy but a hairpin configuration preventing binding to other nanos-tar sticky ends would on average be present 24.2 ±2.1% of the time.

**Figure 5.**
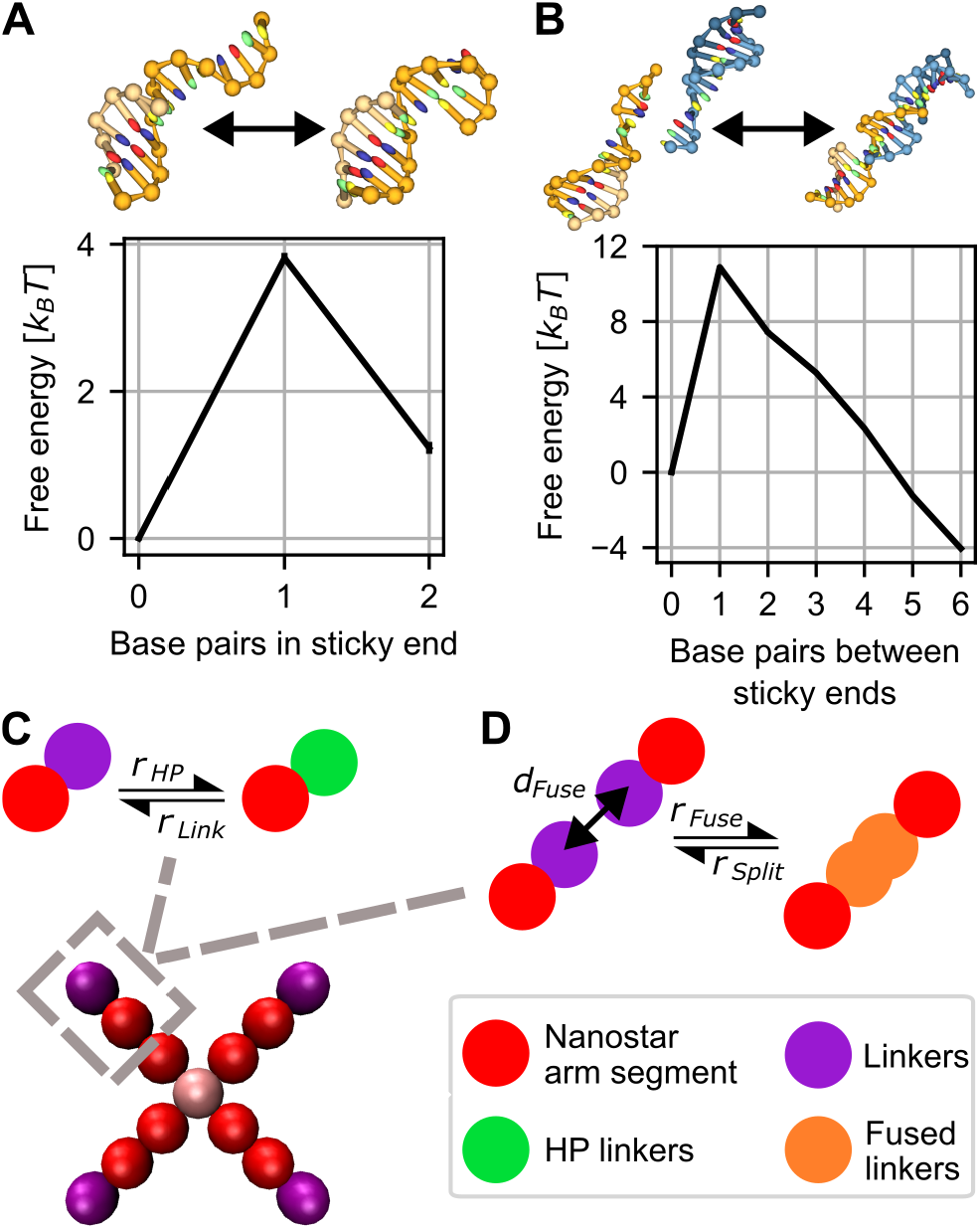
Hybridisation within and between sticky ends. (A) Umbrella sampling was used to obtain free energy profiles of hairpin formation. The simulated structure consisted of the sticky end and a segment of the nanostar arm. The open configuration was favourable. B) Umbrella sampling of hybridisation between sticky ends. The fully hybridised state was most favourable. Data shows mean and standard error of the mean from 6 repeats. (C) In the bead-spring model, a first order reaction with rates *r*_*HP*_ and *r*_*Link*_ toggled the terminal bead between hairpin and free sticky end configuration. (D) A second order reaction fused free sticky ends with rate *r*_*Fuse*_when the distance between their centres was below *d*_*Fuse*_, and split them with rate *r*_*Split*_.

Hairpin formation competes with hybridisation with other sticky ends. Sampling of hybridisation between two sticky ends connected to a short arm segment was carried out with the same strategy used for sampling hairpin formation. Formation of the first base pair was unfavourable due to the large entropic cost and the fully hybridised state was the most stable (Figure 5B). At 20°C, fully or partly hybridised states would thus be present 98.06 ± 0.09% of the time. Bulk correction reduced this value to 90.66 ± 0.23. [57] Note that these percentages are concentration-dependent and may not transfer to the fraction of bound nanostar sticky ends, as the interaction between sticky ends terminating four nanostar arms could be more complex. Nonetheless, the bound fraction obtained for arm fragments indicated that at 20°C the majority of nanostar sticky ends would be fused with other nanostar sticky ends. [39, 74] This finding is also consistent with a prediction of complex formation between sticky ends carried out in NUPACK’s test tube tool. Here, a bound fraction of ≈ 94% was found. Since the sequences tested in NUPACK do not contain attached nanostar arms that constrain the configurational freedom, the bound complex might be more favourable in nanostar networks.

We used the thermodynamic equilibria for hairpin formation and sticky end hybridisation to select rates in our bead-spring model. Hairpin formation in the bead-spring model was implemented as a first order reaction with no spatial dependence (Figure 5C). We selected a ratio between forward and backward rates that produced the same total fraction of time spent in the hairpin formation previously obtained with oxDNA. Hybridisation between nanostar sticky ends was modelled with a second order reaction within a limited distance (Figure 5D). We fixed the binding rate *r*_*Fuse*_ and distance *d*_*Fuse*_ within which sticky ends may fuse and varied the unbinding rate *r*_*Split*_ to obtain partially assembled networks or networks with a fraction of bound nanostars in line with equilibrium predictions from oxDNA and NUPACK.

Based on experimental studies on hybridisation kinetics of short oligos [75] and recent work on similar nanostar systems [40] we estimated that bond lifetimes for the experimental reference design at 20°C are on the order of 100*ms − s*. Since the time scale of diffusive motion simulated in our bead spring model corresponds to the microsecond range, bonds between the sticky ends of the experimental reference should remain largely stable during simulations.

### 3.4 Linking nanostar flexibility and binding affinities to rheological properties

In the following, we used the bead-spring model to assemble nanostar networks, which we interpreted as graphs for extraction of storage and loss moduli. In this context we investigated the influence of sticky end affinities, nanostar flexibility, and hydrodynamic interactions on dynamic moduli.

We placed 100 optimised nanostars in a periodic box under experimental conditions. [39] Each nanostar was treated as a node, with edges formed by sticky-end fusion. After equilibration without reactions, fusion/fission was enabled, leading to gradual formation of a large, connected component (defined here as a set of nanostars linked through hybridised sticky ends). For *r*_*split*_·Δ*t* = 0, 97 ± 0% of sticky ends were bound, and 98.80±0.18 motifs belonged to the largest component (Figure 6A-D). Connectivity approaching the theoretical maximum is expected under the conditions of the experimental reference, where a bound fraction around 98% was suggested. [39] The resulting G’ and G” curves (Figure 6C) appeared near-Maxwellian and lacked a pronounced intermediate scaling regime.

**Figure 6.**
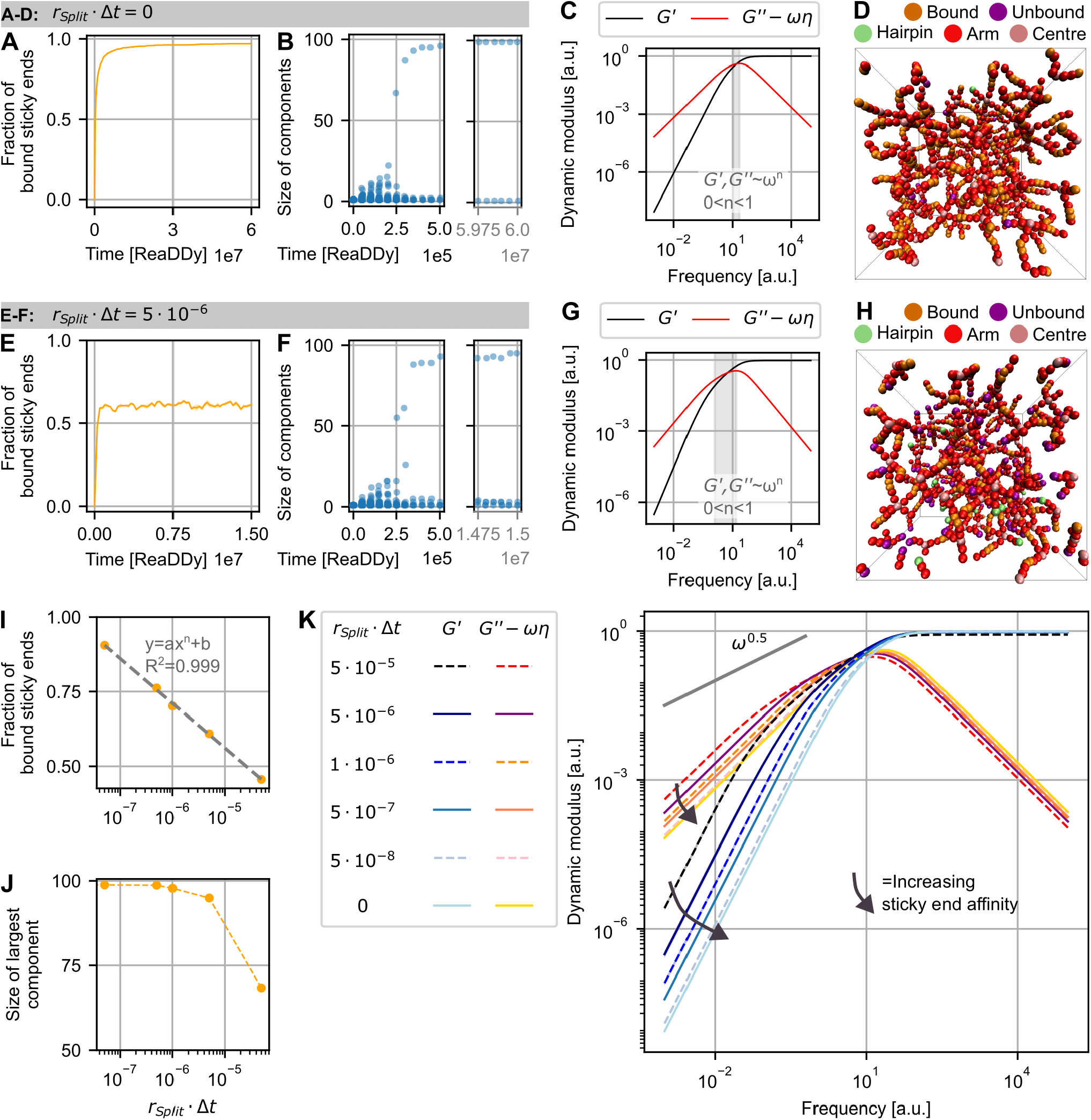
Rheological properties depend on the level of networking. (A) For *r*_*Split*_ · Δ*t* = 0.0, the fraction of bound (hybridised) sticky ends approached one. (B) Each dot in the stripplot corresponds to 1 component in a simulation. Initially, multiple small components were formed which fused into larger components until all but one nanostar were part of one component. (C) With almost all sticky ends hybridised, the intermediate scaling regime (grey) for G’ and G” mostly vanished. (D) Snapshot at time step 1.5 · 10^7^, most sticky ends were bound. (E) For *r*_*Split*_ · Δ*t* = 5 · 10^*−*6^, *≈* 61% of all sticky ends were bound. (F) A dominant component emerged in the simulation, but multiple smaller components remained separate. (G) G’ and G” now featured a pronounced intermediate scaling regime (grey). (H) Snapshot at time step 1.5 · 10^7^, the network contained notably more unbound sticky ends. (I) Logarithmic dependence of fraction of bound sticky ends on splitting rate. (J) Decrease in the number of nanostars within the largest component with increasing splitting rate. (K) Changes in predicted G’ and G” curves for increasing connectivity between sticky ends. Figure only shows changes in scaling, without rescaling to account for expected changes in absolute values. Fraction of bound sticky ends, largest components and dynamic moduli show mean and standard error of the mean (often within line width) averaged from five simulations of 100 nanostars.

To probe lower connectivity, we set *r*_*split*_ · Δ*t* = 5 · 10^*−*6^, yielding 60.9 ±0.5% bound sticky ends and a largest component containing 94.93±0.24 motifs (Figure 6E-H). We observed a heterogeneous valence distribution within these networks, meaning that some nanostars were disconnected, or connected with only 1, 2 or 3 sticky ends. Over time, the network reconfigured through splitting and fusion events, fluctuating around a stable mean fraction of sticky ends. Here, G’ and G” (averaged over many network configurations) exhibited a clearer intermediate scaling regime (Figure 6G). Such a scenario could correspond to a partially assembled network (e.g., during a gradual cool-down) or be achieved by introducing weaker base pairs, mismatches in the sticky ends, making the sticky ends shorter or increasing the temperature. To give concrete examples, we used our Bayesian optimisation strategy to predict a library of sticky end sequences, temperature and salt conditions in line with a bound fraction of ≈ 61%, which can be found in the SI (Table S2).

Simulations of network formation using our bead-spring model required only a few CPU hours per design, enabling rapid screening of sticky-end affinities. We included results from five additional affinities, producing fractions of bound sticky ends that decreased logarithmically with increasing splitting rate, reaching a minimum of 45.6 ± 0.4% (Figure 6I). At low splitting rates, the size of the largest component converged to 100 because at any given time most nanostars were hybridised via their sticky ends to form a single, system-spanning network (Figure 6J). With lowered fractions of bound sticky ends, G’ and G” curves revealed a continuous shift from negligible intermediate scaling range to a pronounced frequency range where the magnitudes of moduli run close to each other and scale G’, G” ∼ *ω*^*n*^ with 0 *< n <* 1 (Figure 6K). For simulations with fractions of bound sticky ends approaching the percolation threshold of 1/3 according to the Flory-Stockmayer criterion [76, 77], G’ and G”curves fluctuated, indicating that extended networks were needed to sample relaxation times for consistent predictions.

We wondered if the affinity-dependent networking would also be sensitive to nanostar flexibility. To test this, we set the spring constants of the inter-arm potentials (*k*_*base*_, *k*_*oppa*_, *k*_*bound*_) near zero, resulting in broader angle distributions (Figure 7C). With this choice we wanted to assess if the structural rigidity provided solely by angle potentials within arms and repulsion between beads would result in similar networking and viscoelasticity. Simulations without inter-arm potentials showed slightly higher bound sticky end fractions but smaller largest components, indicating that flexibility promoted hybridisation but reduced network connectivity (Figure 7A,B). However, for both rates *r*_*Split*_ · Δ*t* = 5 · 10^*−*6^ and *r*_*Split*_ · Δ*t* = 0 the differences between fully parametrised and no inter-arm potential were minor (Table 1). Without inter-arm potentials, G’ and G” shifted upward at low frequencies and displayed extended intermediate scaling (Figure 7E,F). Networks without inter-arm potentials appeared less ordered in simulations snapshots and exhibited local density variations (Figure 7D). Radial distribution functions obtained from motif centres confirmed reduced ordering: peaks broadened and shifted to smaller radii (Figure 8).

**Table 1.**
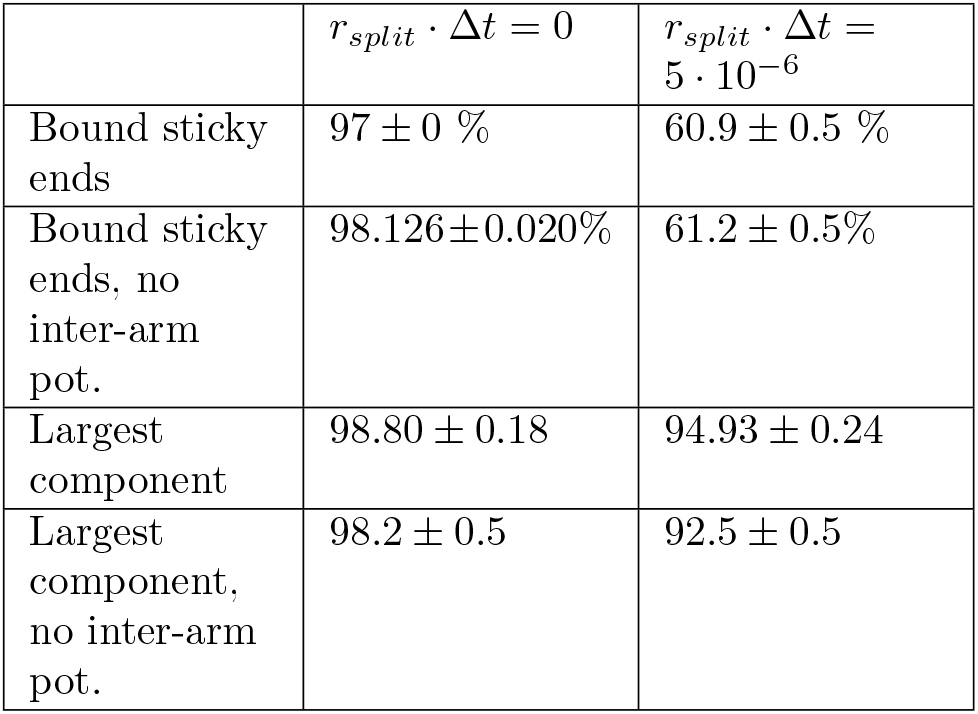
Bound sticky end fractions and largest component sizes for different sticky end affinities and local mechanical flexibility. Mean and standard error of the mean obtained from five repeats.

**Figure 7.**
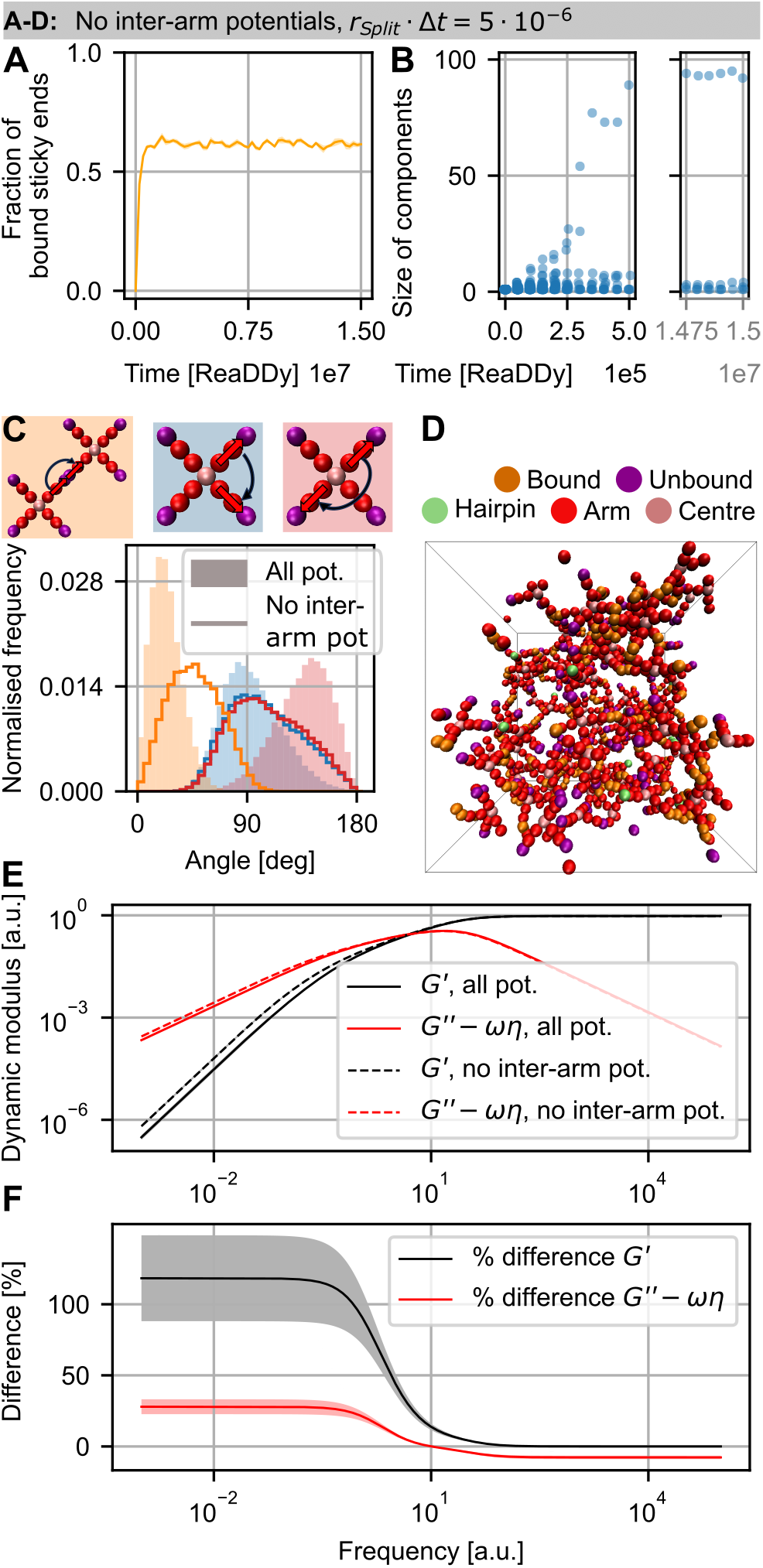
Rheological properties depend on the inter-arm nanostar flexibility. (A), (B) Using *r*_*Split*_ ·Δ*t* = 5·10^*−*6^ but setting the inter-arm angle potentials to a value close to zero did not significantly influence the fraction of bound sticky ends or the size of the largest component in comparison to the case with inter-arm potentials seen in Figure 6 E,F. (C) Broader angle distributions (plotted non-filled) as consequence of near zero inter-arm potentials. (D) Increased inter-arm flexibility lead to variations in nanostar densities, as seen in a snapshot at timestep 1.5 · 10^7^. (E) G’ and G” had a stretched intermediate scaling regime without inter-arm potentials, leading to up-shifted dynamic moduli at low frequencies. (F) Percentage difference between moduli in E, normalised to the moduli with inter-arm potentials. Fraction of bound sticky ends and angle potentials averaged from five repeats. Curves in A, E and F show mean and standard error of the mean from five repeats.

**Figure 8.**
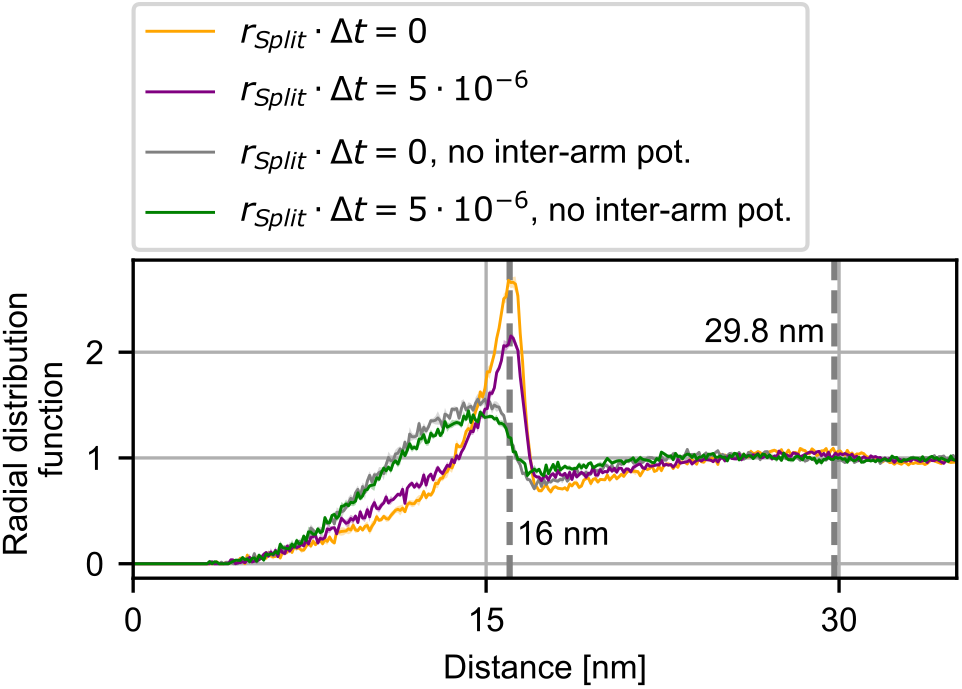
Radial distribution function in nanostar networks. Radial distribution functions from simulations with and without inter-arm potentials for high (*r*_*split*_ · Δ*t* = 0) and low (*r*_*split*_ · Δ*t* = 5 · 10^*−*6^) nanostar connectivity. Peak positions obtained from simulations with inter-arm potentials. Curves show mean and standard error of the mean from 51 configurations of the final 3.75 · 10^6^ time steps from five repeats.

### 3.5 Hydrodynamic interactions narrow relaxation spectra and reduce moduli away from cross over

To include hydrodynamic interactions (HI), we modified the calculation of relaxation times and moduli by pre-averaging the mobility matrix (**Ĥ**) and multiplying it by the graph Laplacian 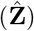. This mobility matrix introduces long-range spatial couplings between all nanostars based on the inverse equilibrium distances between nanostars and an effective hydrodynamic radius. We estimated that this effective radius is approximately *a* = 4.7 nm for each 4-armed nanostar based on experimental measurements, [17] and note that slightly smaller or larger values did not significantly change the results. The distance-depended couplings between nanostars were calculated by averaging the inverse centre-centre distances in each equilibrated network. We thus approximated the hydrodynamic interaction as a spherically symmetric coupling centred on each nanostar’s centre. We observed a compression of the relaxation time spectrum upon the inclusion of HI (Figure 9A). This compression could be explained by differential speed-up across modes: while fast modes are enhanced by local hydrodynamic lubrication, slow modes experience a more pronounced acceleration due to cooperative solvent motion and partial shielding, reducing the effective friction on the chain. While the first mode in Figure 9A for *r*_*Split*_·Δ*t* = 0 was sped up by ≈ 1.6%, last mode was sped up by ≈ 39%. This narrowing of the spectrum is consistent with the theoretical shift in relaxation time scaling from *τ*_*p*_ ∝ *p*^*−*2^ in the free-draining Rouse model to *τ*_*p*_ ∝ *p*^*−*1.5^ in the non-free-draining Zimm model (SI Section S6 and Figure S8). Figure 9B illustrates the storage and loss moduli for simulations at varying *r*_*split*_·Δ*t*, including the solvent contribution to *G*^*′′*^. This highlights the transition from intermediate scaling (approaching *ω*^1*/*2^) to a high-frequency regime dominated by the purely viscous solvent scaling (*ω*^1^). When comparing the results with and without HI (Figure 9C,D), a systematic shift in the moduli was observed. Upon aligning the crossover points, the inclusion of HI results in a predominantly negative shift, with differences reaching approximately 24% in the low-frequency limit.

**Figure 9.**
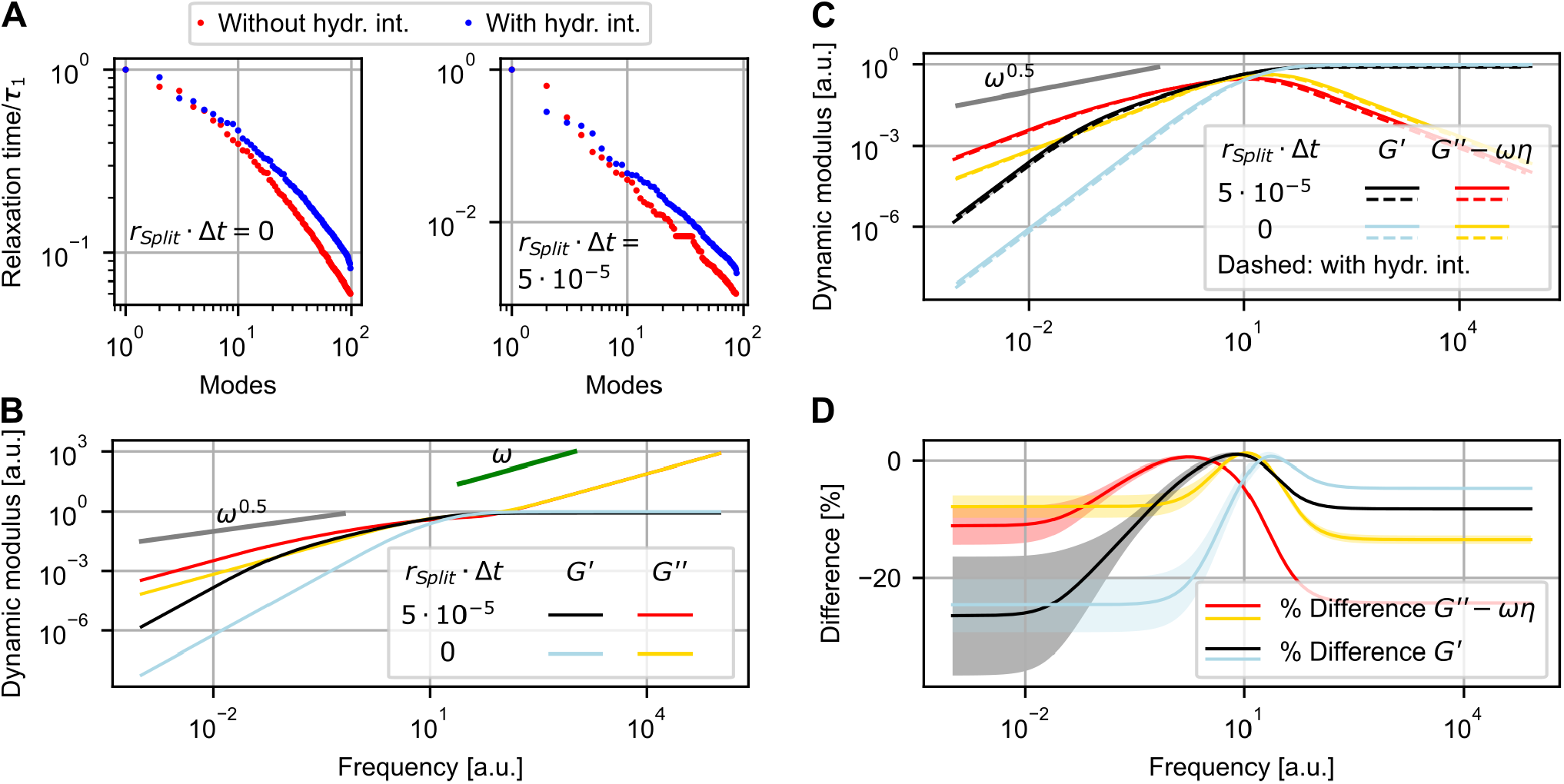
Influence of hydrodynamic interactions on relaxation times and moduli. (A) Including the pre-averaged Oseen tensor to approximate hydrodynamic interactions between nanostars compressed the observed relaxation spectra. For comparability, all relaxation times were normalised by the first relaxation time. (B) Moduli extracted with inclusion of hydrodynamic interaction. Shown with the solvent contribution to the loss modulus *ωη*_*solv*_, with *η*_*solv*_ = 0.008 [a.u.] for illustration. (C) Predicted moduli with (dashed lines) and without hydrodynamic interactions (solid lines). (D) Difference between moduli without and with inclusion of hydrodynamic interactions. Relaxation times extracted from one simulation for each condition. Mean and standard error of the mean for moduli from five repeats each.

### 3.6 Reproducing experimental bulk-rheology required relaxation times from optimised simulations

Building on the flexibility- and affinity-dependent rheological responses, we asked if relaxation times from optimised simulations could reproduce experimental data.

The near-Maxwellian scaling for high sticky end connectivity qualitatively matched experiments well below the sticky end melting temperature. [11, 41] Closer to the melting temperature, intermediate scaling approaching *∼ω*^1*/*2^ with small separation between G’ and G” has been observed, consistent with our results for reduced connectivity between sticky ends. [39, 40, 42] We directly compared our results to experimental data for the same design and conditions used in our simulations by introducing physical units. To this end, following recent work, [43, 73] we used a rescaling that multiplied frequency and moduli with constants to match the cross over between G’ and G” to that of the experiment.

Moduli extracted from highly connected simulations (*r*_*split*_ ·Δ*t* = 0) aligned closely with experimental moduli (Figure 10A). Underestimation of measured G” at high frequencies could be caused by multiple sources: The graph-based method produced G” values dereferenced against the medium viscosity while the experiment recorded such hydrodynamic effects. Furthermore, the precision of the experiment is limited at high frequencies, for instance by motor inertia, which can lead to an artificial upshift in G”. [78]

**Figure 10.**
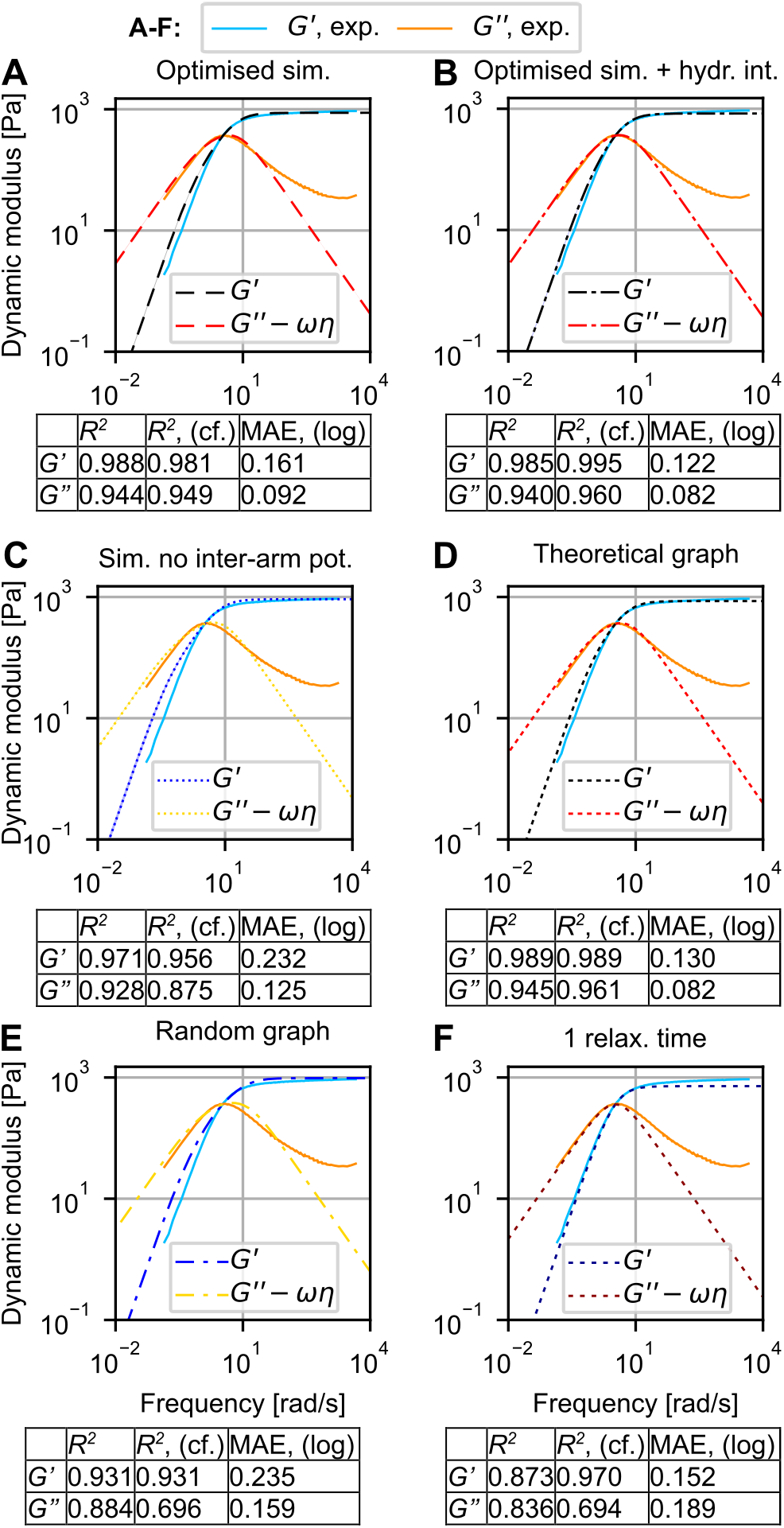
Comparing dynamic moduli from experiment, simulation and theory. (A) G’ and G” extracted from graph interpretation of optimised nanostar simulations and from experiments. (B) Moduli from simulations without inter-arm potential. (C) G’ and G” extracted from a theoretical graph representation of a fully connected network of four-armed molecules (nodes). (D) Moduli from random graph generated with maximum node degree of 99. (E) Approximation of G’ and G” with a Maxwell model with one relaxation time. G’ and G” curves from theory and simulation were rescaled to match the experimental crossover frequency. Curves from simulations averaged five repeats. *R*^2^ values were calculated for the full experimental frequency range or constrained to 0.5-50 rad/s (indicated by cf.). The mean absolute error (MAE, (log)) integrated the absolute error in log-space. MAE values for G” where only integrated up to 50 rad/s, excluding the region with systematic deviations.

To quantify how well the graph-based predictions match the experimental data, we calculated *R*^2^ values across the full experimental frequency range and in a region around the crossover frequency (0.5-50 rad/s). We also computed the mean absolute error (MAE) via integration in log space to quantify the deviation apparent in the commonly used double logarithmic plots. These metrics indicated that the fully optimised model produced a closer match to the experimental moduli than the model without inter-arm potentials (Figure 10A,C). The better fit of the fully parametrised model highlights the complex interplay between local flexibility, emergent network structure and viscoelastic behaviour, underscoring the relevance sequence-informed coarse graining. We previously observed that inclusion of HI lowered the predicted moduli, especially for low frequencies. Indeed, the slight overestimation of experimental moduli for low frequencies was notably reduced when we included HI, indicating that inclusion of the pre-averaged mobility matrix can improve predictions, especially for low frequencies. (Figure 10B)

We compared the predictions made from the graph-based method, which used a spectrum of relaxation times, to the predictions made by a conventional Maxwell model with only one relaxation time. In this baseline model the relaxation time is the inverse of the crossover frequency, calculated from the experimental data. Using a single relaxation time to generate G’ and G” produced the observed scaling regimes and matched the experimen-tal values well for frequencies below the crossover. Above the crossover the predictions deviated notably and the plateau modulus was underestimated (Figure 10F). For example, at a frequency of 63 rad/s, used in [39] to assess the plateau modulus, G’ measured ≈826 Pa. Extraction from optimised simulations predicted 869 Pa, additional inclusion of HI yielded 830 Pa while using a single relaxation time predicted 718 Pa (Figure 10A,B,F).

The graph-based method applied to the optimised simulations with HI outperformed the conventional Maxwell model across all metrics. Without HI, the graph based method performed better than the conventional model in all metrics with the exception of the MAE value for G’. This finding reflected the fact that in log-space, the conventional Maxwell model produced a slightly better approximation of G’ for low frequencies, which offset the deviations at high frequencies (Figure 10A,E). We also note that the graph-based model with HI performed better than the model without HI around the crossover frequency and in log space, but slightly worse when considering the full frequency space. The slight reduction is caused by the dominant high frequency contribution far from the crossover, which was marginally better approximated without HI.

### 3.7 Theoretical graphs of nanostar materials predict rheological behaviour

Optimised simulations revealed highly connected, homogeneous nanostar networks. We asked whether generating graphs of such networks theoretically could reproduce experimental moduli and enable efficient predictions.

Graphs were generated by specifying maximum node degree (i.e., motif valency) and edge formation probability (i.e., affinity between motifs). A random graph with degree four (representing four-armed nanostars below the sticky end melting temperature) produced G’ and G” curves shown in Figure 10C. Comparison to the experimental data yielded *R*^2^ and MAE values that marginally improved upon the results extracted from optimised simulations without HI. The slightly higher connectivity of the theoretical graph improved the predictions in the low frequency range, and outperformed the conventional Maxwell model across all metrics (Figure 10A,C,E). The predictions from theoretical graphs performed similar to the predictions from optimised simulations with HI. This apparent similarity could indicate that the perfect connectivity of the theoretical graph, which was not achieved in simulations, could offset neglecting additional coupling due to hydrodynamic interactions. An alternative interpretation would be that the experimental connectivity is indeed slightly higher than in simulations, potentially because of limited simulation time and limited possibility to reorder unfavourable sticky end hybridisation with *r*_*split*_ · Δ*t* = 0.

To test the sensitivity of theoretical graphs to molec-ular organisation, we kept node and edge counts (100 nodes, 400 edges) but allowed a maximum degree of 99. Resulting G’ and G” curves varied strongly between random graphs and deviated from experiments (Figure 10D), confirming that valency and edge probabilities critically shape rheology.

We extended the analysis to 3-, 5-, and 6-armed nanostars. Graphs constrained to the correct valency produced G’ and G” curves with similar scaling that matched experimental data [39] well for 5- and 6-armed motifs (Figure 11A). For 3-armed motifs, deviations appeared above and below the crossover frequency (Figure 11C). We investigated whether simulations of networks from 3-armed nanostars would produce a better fit than prediction from theoretical graphs. (SI Section S6 and Figure S9) To this end, we utilised a bead-spring model that we optimised for a different project using a slightly different sequence design and conditions, and applied the conditions of the experimental reference. Predictions were improved, especially for high frequencies, however, the predictions both with and without HI could not fully account for the deviations. The better agreement between theoretical predictions and experimental curves for nanostars with higher valency might stem from the generalised Maxwell model being more applicable near the isostatic point, where nodes are constrained and linker stretching dominates. In contrast, bending contributions become significant in underconstrained networks, such as those formed by three-armed nanostars [39]. Interestingly, the 3-arm data was better captured either by increasing network connectivity with nodes of higher degree or by limiting the relaxation time spectrum. Both approaches effectively constrain the range of allowed relaxation times. This suggests that 3-armed motifs may be more constrained than expected from connectivity alone, possibly due to topological linking, entanglements, or excluded volume effects. Topological linking, in particular, was identified as contribution in networks of three-armed nanostars at high concentration. [40]

**Figure 11.**
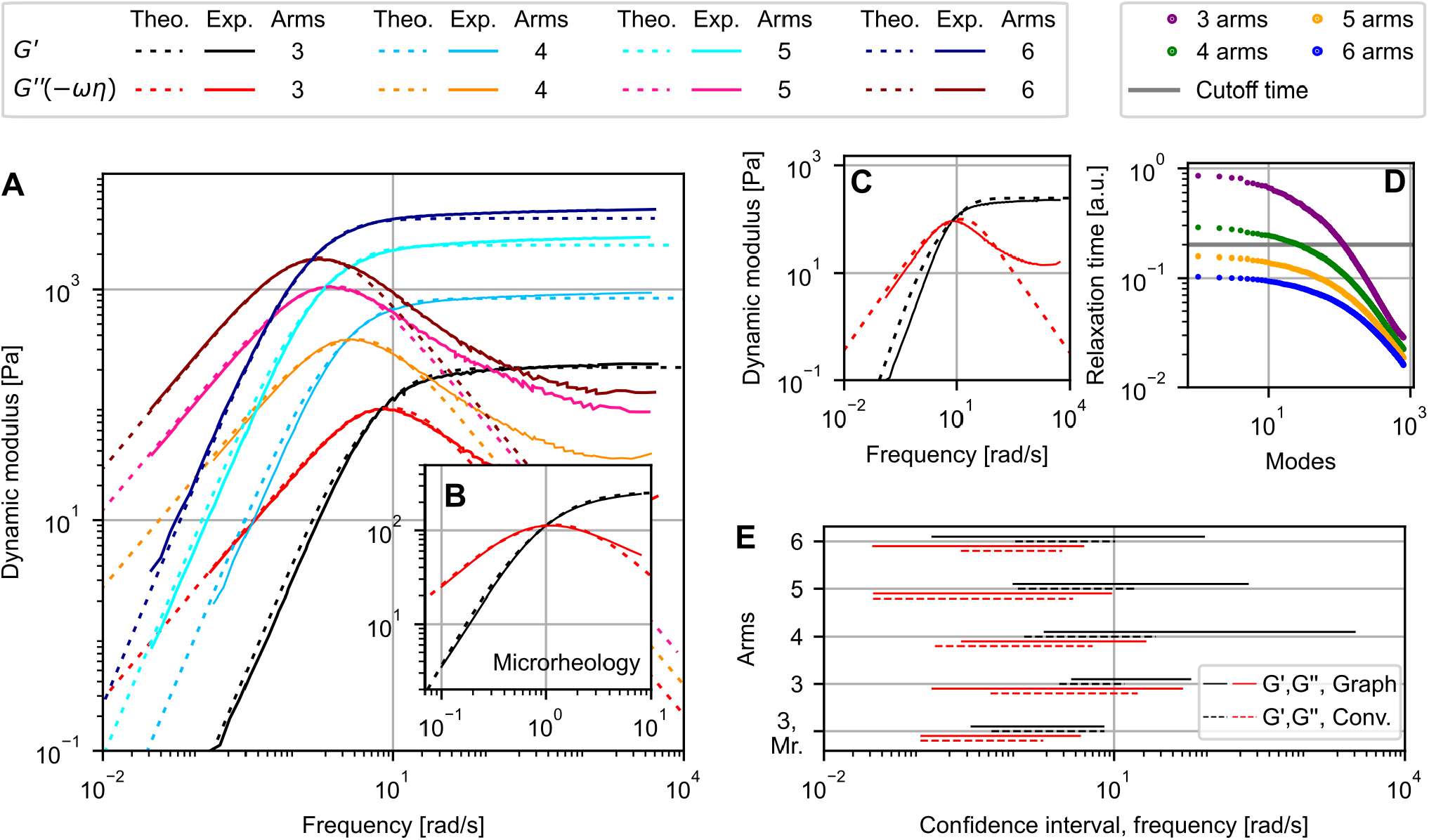
Comparing theoretical and experimental rheology for networks of nanostars with different numbers of arms. (A) Rescaled predictions for G’ and G” extracted from networks with node degrees 3, 4, 5 and 6 compared to experimental bulk rheology data [39] for nanostar networks with 3, 4, 5 and 6 arms. Relaxation spectra for nodes with degrees 3 and 4 were constrained by a cutoff given in D. (B) Comparison of theoretical results to microrheology data for 3-armed nanostars. [40] (C) Predictions for 3-armed nanostars with full relaxation spectrum. (D) Relaxation time spectra with cutoff. (E) Contiguous confidence regions in which predictions from the graph based and a conventional Maxwell model deviate *≤*10% from experiments. Theoretical graphs used 800 nodes.

Phenomenologically, introducing a cutoff in the relaxation time spectrum positioned between the longest relaxation times of degree-four and degree-five networks, improved predictions for the 3-arm motif and marginally for the 4-arm motif. (Figure 11A,D). Further, predictions aligned well with microrheology data for similar 3-arm designs (Figure 11B), [40] suggesting that predictions are not limited to bulk rheology. Consistent with our observations for 4-armed nanostars, *R*^2^ and MAE values showed that a spectrum of relaxation times approximated the experimental data more accurately than a single relaxation time (SI, Table S3). To visually represent frequency regions in which either method makes accurate predictions, we plotted contiguous frequency intervals where predictions for G’ and G” deviated ≤10% from experimental moduli. Graph-based methods produced wider frequency ranges where predictions closely matched experiments. The better approximation of G’ and G” by using a spectrum of relaxation time was most notable for high frequencies corresponding to the plateau modulus (Figure 11E).

We also tested whether the total number of nodes influenced the predictions for G’ and G” but found only marginal influence when using between 100 and 800 nodes. For much lower node counts (10-20), we found up-shifted moduli and deviations, particularly for degree 5 and 6 networks, indicating insufficient sampling from the relaxation time spectrum. In the conventional Maxwell model, the crossover frequency is the inverse of the single relaxation time: 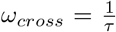 . Empirically, we found this scaling to be conserved when applying the generalised Maxwell model to networks with increasing connectivity and evaluating the inverse of the mean relaxation time. Observed crossover frequencies scattered around a function 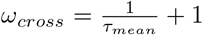 with *R*^2^ values of 0.99 for 1000 networks with 100 nodes of maximum degree 4 and between 50 and 400 randomly added edges. Increasing the maximum node degree to 99 produced similar scaling with *R*^2^ values of 0.95.

### 3.8 Optimising sticky ends to match desired rheology

The methods demonstrated thus far can be combined into workflows for inverse design of viscoelastic DNA materials. In the following we showcase two approaches that take a target viscoelastic response and predict the required sticky end sequences and conditions for corresponding materials.

Approach one used the viscoelastic response and fraction of bound sticky ends of five optimised simulations as training and test data. The five simulations differed in the splitting rates between sticky ends, modelling different hypothetical affinities. G’ and G” were extracted according to the graph-based, generalised Maxwell model, without inclusion of HI for simplicity. One simulation served as a hypothetical optimisation target, the other four, covering bound fractions of sticky ends between ≈47% and ≈90%, were used for training. A Gaussian process model predicted what fraction of bound sticky ends would maximise similarity to the target G’ and G” curves. In the example shown in Figure 12, this approach predicted the fraction of bound sticky ends (≈71%) within the standard error of the mean, without requiring further rounds of optimisation. This fraction was then converted into candidate sequences via Bayesian optimisation coupled to NUPACK’s test tube tool. Optimisation (100 initial designs, 300 iterations) produced dozens of sequences, salt and temperature conditions that matched the target fraction of bound sticky ends within minutes of computation time. Exemplary sticky end sequences with predicted bound fractions 71 ± 1% (Figure 12) were chosen from parameter ranges close to the optimised simulations, ensuring the validity of the material’s network structures. Additional simulations optimised for other parameter regions could be used for further fine-tuning steps or validation. This approach benefits from physically meaningful simulations that capture mechanical and thermodynamic properties of DNA.

**Figure 12.**
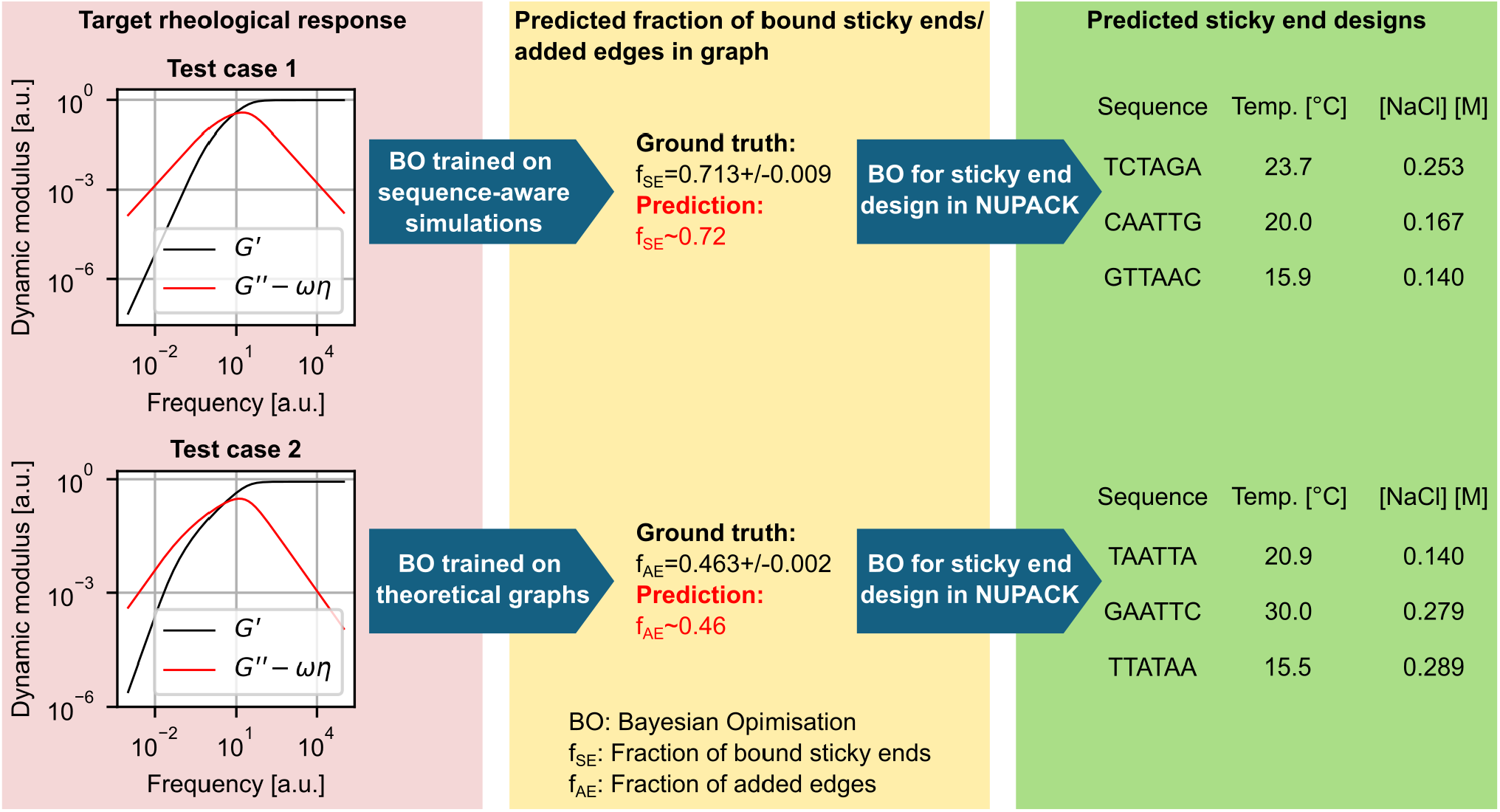
Inverse design of sticky end sequences to match target rheology. Bayesian optimisation trained on network connectivity and resulting G’ and G” curves can predict connectivity required for target G’ and G” curves. For illustration, the two target cases were obtained from simulations (averaged from five repeats each). Case 1 shows prediction using 4 optimised simulations for training on physically realistic networks. Case 2 used 86 theoretical graphs for training, trading network realism for more training data. The unknown ground truth was accurately predicted in both cases. Bayesian optimisation of free energies can then predict corresponding sticky end sequences, temperature and salt conditions.

If the underlying graph structures are well understood through optimised simulations and comparison to experiments, one can use theoretical graphs which are much faster to generate. For the second approach we replaced the simulations with theoretical graphs of degree 4 and selected 86 fractions of realised edges (i.e., hybridised sticky ends), covering the range from percolation to maximum connectivity. For each fraction we averaged the viscoelastic response of 10 random graphs. By using the theoretical graphs as training data, the fraction of added edges of the five test simulation used for the first approach was predicted within the standard error of the mean, when G’ and G” curves were used as targets. We assumed that this fraction of added edges is equivalent to the required fraction of bound sticky ends, which allowed to proceed with the sequence prediction step using NU-PACK as outlined before. Depending on the flexibility of nanostars, arms of the same nanostar might be fused, or two nanostars might be fused via more than one pair of arms. For the tested design this was rarely the case and the difference between fractions of fused sticky ends and edges in the graph was less than 1%. For more flexible designs, graphs with self-loops or multiple edged between nanostars should be considered. Successful prediction of the underlying sticky end fraction (≈46%) for a second test case where the desired viscoelastic response featured a pronounced intermediate scaling regime can be seen in Figure 12.

Both approaches, while applicable to simulated data, likely contain remaining uncertainties when applying them to experiments. First, comparison to a larger set of experiments will determine under which conditions and limitations the generalised Maxwell model is applicable. Second, NUPACK neglects constraints from sticky end attachment to nanostar arms. This (de-)stabilisation could be corrected with an additive free-energy term.

## 4 Discussion

Our work develops and applies a computational framework that links molecular design to viscoelastic behaviour in cross-linked polymer networks. Methodologically, we link simulations at different scales with Bayesian optimisation, enabling scalable modelling without reliance on molecule-specific details. At the mesoscopic level, we interpret cross-linked polymer materials as graph-based generalisations of classical Rouse and Zimm models. This perspective allows us to predict relaxation-time spectra and the scaling of storage and loss moduli directly from simulated or theoretical network topologies.

We apply this framework to parametrise a DNA-sequence-informed bead-spring model of DNA nanostars. The connection to underlying sequences was established by reproducing structural flexibility recorded with nucleotide-level simulations. Harnessing increased computational efficiency, the bead-spring model was then used to investigate the assembly of DNA nanostars into networked materials. To investigate rheological properties, we represented DNA networks as graphs, where nodes correspond to nanostars, and edges to hybridised sticky ends. This approach enables efficient exploration of the design space, connecting DNA sequences with nanostar flexibility, sticky end affinity, and the viscoelastic properties of the resulting networks. The independence from system specific details could make Bayesian optimisation and graph-analysis broadly applicable by linking tuneable material parameters to desired structural or thermodynamic properties.

In hydrogel networks of DNA nanostars, simulations revealed how sticky end affinity and nanostar flexibility influence connectivity and resulting graph structures. By analysing the eigenvalues of the graph Laplacian, we extracted relaxation time spectra for a generalised Maxwell model. [43, 73] By multiplying the graph Laplacian with the pre-averaged mobility matrix, we demonstrate how hydrodynamic interactions can be accounted for in a graph-based interpretation of the Zimm model. Including hydrodynamic interactions lead to a compressed relaxation time spectrum, which we interpreted as a differential speed up of relaxation processes due to hydrodynamic interactions. The graph-based approach was sensitive to molecular architecture, captured through node degree and connectivity. For lower connectivity, the graph-based approach predicted the appearance of a growing region where G’, G” ∼*ω*^*n*^ with 0 *< n <* 1, approaching *n* = 0.5, and with shrinking separation between magnitudes of G’ and G”. A frequency region with moduli scaling proportional to *ω*^0.5^ with similar magnitudes has been experimentally observed close to the melting temperature, [11, 41, 42] and links to the gel point of a material. [79] At the gel point, often associated with a divergence of the longest relaxation time, elastic and viscous contributions are comparable, and an infinite, percolating cluster appears. [80] A link between relaxation time and connectivity is in line with the graph-interpretation: The smallest non-zero eigenvalue of the graph Laplacian of a connected graph quantifies the connectivity. [81, 82] Thus, lowering connectivity towards percolation increases the longest relaxation time. While our predictions captured observed changes in scaling, the high frequency plateau modulus is often absent in experiments at higher temperatures (50-60°C).[11, 41, 42] At these temperatures, bond lifetimes may be too short for a plateau to appear, indicating limitations in extending the assumed relaxation spectrum to elevated temperatures. Simulations of optimised four-armed nanostars at 20°C revealed a graph with highly connected nodes of degree four, motivating a theoretical extension to nanostars with varying arm numbers. Evaluating such graph-based material representations provided robust predictions for viscoelastic behaviour. Experimental scaling was captured and predictions closely matched experimental moduli [39, 40] after applying a simple rescaling transformation. Compared to the conventional single-mode Maxwell model, incorporating a spectrum of relaxation times improved predictions particularly near and above the crossover frequency, extending to the plateau modulus. Predictions where further refined by including hydrodynamic interactions, which lowered predicted moduli for low and high frequencies without changing the observed scaling, leading to a better prediction for all but the highest frequencies.

Combining these scale-bridging methods into a modular workflow, we showcased inverse design of viscoelastic DNA materials. Starting from desired rheological responses, we predicted underlying network connectivity which we translated into sticky end sequences, salt and temperature conditions. These inverse design strategies will benefit from integration of future experimental studies that will inform more accurate theoretical descriptions, but can already define and demarcate interesting design sub-spaces for experimental investigations.

Our results place DNA nanostar hydrogels within the broader class of transient, limited-valence soft materials whose linear rheology is governed by the interplay of bond lifetime, network connectivity, and topology. The graph-derived relaxation spectrum used here provides a structural interpretation of the emergence of different rheological responses. Our approach is complementary to fractional Maxwell descriptions, which capture similar broad-spectrum relaxation phenomenologically but do not directly resolve the underlying architecture. We note that a fractional Maxwell model has recently been applied to model the viscoelastic response of DNA nanostar hydrogel in a study that also reported solid-like behaviour at high temperatures well above the percolation threshold, which was attributed to a glass transition. [46] A different method to incorporate hydrodynamic interaction was recently proposed for biomolecular condensates. [83] Here, networks of Maxwell elements in a Stokes fluid allowed to carry out creep test and to characterise viscoelastic moduli. Exploring different values of elasticity and viscosity produced a range of scaling behaviours, including Maxwell-like, as well as Rouse and Zimm regimes, providing insights into the parameter selection required to match experimental scaling.

The emergence of power-law scaling in our simulations mirrors recent findings in patchy particle systems. [48] Evaluation of the stress autocorrelation in simulations showed that the addition of divalent particles to networks of tri- and tetravalent particles induced a departure from terminal Maxwellian behaviour. For an increasing fraction of divalent patches, the scaling exponent for G’ approached ≈ 0.6. [48] In the DNA nanostar networks simulated here, lower effective sticky-end affinities similarly resulted in a distribution of transiently unbound arms, suggesting that valence polydispersity could by a universal driver for the observed *ω*^*n*^ scaling. Power law scaling has also been reported for fractal colloidal gels, [49] where the linear rheology was described by a fractional Maxwell model and was linked to reversible non-affine rearrangements in a connected, fractal network. These findings suggest that power-law rheology can arise from heterogeneous relaxation of complex network topologies.

A connection between polymer cross-linking and changes in observed rheology was also reported for hydrogen-bonded supramolecular networks, where adding comonomers that more strongly cross-linked the polymers shifted the observed scaling of storage and loss modulus from near-Maxwellian at low frequencies towards *ω*^*n*^, *n* = 0.5 *−* 0.7 with a notable plateau modulus. [84] Similarly, the rheology of hydrophobically modified ethoxylated urethane (HEUR) thickeners transitioned from single-mode to generalised Maxwell behaviour by the formation of a transient, percolated network of bridged latex particles. [85]

Conceptually, these findings resonate with recent models of multivalent reversible gels, in which increased cross-linker valence generates more heterogeneous bonding pathways and broadens the distribution of relaxation timescales, further demonstrating how network architecture governs rheology. [86]

Our integration of machine learning, simulations and graph theory reflects a broader move toward predictive material design in (soft) matter science. Related approaches have recently been used to automate the exploration of complex structural spaces, for example by applying active learning to identify polymer architectures with targeted rheology [87] or by using multi-agent frameworks [88] and variational autoencoders [89] to link polymer topology to material properties. Furthermore, using optimised coarse-grained simulations and the graph Laplacian for rapid evaluation mirrors the development of computational high-throughput screening tools for complex bio-architectures, including helical fibrils [90] and polypeptide-based compartments. [91]

Looking ahead, machine-learning-guided optimisation of force fields for general-purpose simulation platforms such as ReaDDy could accelerate the design of functional DNA-based materials. These materials could be hybrid systems containing other types of molecules [12, 13, 92] and non-equilibrium processes such as chemical reactions.[22, 93] Computational workflows could complement experiments for precise control over phase behaviour, [17, 20, 23] mechanical properties,[11, 27, 41, 41, 94, 95] and environmental responsiveness. [6, 14] Controlling such material properties could enable applications in biomimetic systems, [9, 10] and DNA-based mechanical or biochemical computing. [6–8]

Finally, graph-based rheology can complement molecular simulations by bridging molecular architecture and emergent rheology. A current limitation is that theoretical prefactors in the generalised Maxwell model were set to unity. To predict absolute values for frequencies and moduli, future work could derive values of prefactors for DNA nanostars or establish empirical rescaling methods that are transferable to other designs. Another avenue would be to use a relationship between topological linking and the value of the plateau modulus that was recently proposed. [40] The fact that graph-based analysis can provide insights into scaling of moduli with frequency and into scaling of the plateau modulus with concentration, [40] hints at close connection between viscoelasticity and networking that is not fully understood yet. Future extension of graph-based methods could combine connectivity analysis with topological information, for instance by considering loops created by nanostars, which effectively form a network-within-network structure. [40] Simulations that link molecular to material scales seem particularly suitable to deepen our understanding of how topology, connectivity, and rheology interact across scales.

## Author contributions

AG: Conceptualisation (Equal), Formal analysis (Lead), Investigation (Lead), Methodology (Lead), Software (Lead), Validation (Lead), Visualisation (Lead), Writing - original draft (Lead), Writing - review & editing (Equal) LH: Conceptualisation (Equal), Funding acquisition (Lead), Resources (Lead), Supervision (Lead), Writing - review & editing (Equal)

## Data availability

The code to run and evaluate Bayesian optimisation, oxDNA and ReaDDy simulations is available on GitHub (https://github.com/aaron-gad/DNA_Nanomotif_ML_MD). Code and data generated for this work are archived on Zenodo (https://doi.org/10.5281/zenodo.20022811).

## Acknowledgements

We thank Nathaniel Conrad for sharing experimental data on G’ and G”, and for engaging in insightful discussions alongside Deborah Fygenson and Omar Saleh regarding the rheological results and their comparison to experimental observations. We thank Pascal Friederich for helpful discussions regarding the use of Bayesian optimisation and the group of Moritz Kreysing for access to compute resources. We thank Xenia Schneider for additional simulation work on 3-armed nanostars. Large language models (Copilot, Perplexity, Gemini) have been used to assist in drafting text and generating code snippets during the preparation of this manuscript. All outputs were reviewed and validated by the authors. This work is supported by the Helmholtz Association program Natural, Artificial and Cognitive Information Processing, the Helmholtz Initiative and Networking Fund on the HAICORE@KIT partition and by funding from the Carl-Zeiss-Stiftung via the Center SynGen. None of the funding sources have specific grant numbers assigned. This research was edited by our textician, Daniel Shea.

## References

[1] Seeman, N. C. Nucleic acid junctions and lattices. Journal of Theoretical Biology 99, 237–247 (1982).

[2] Rothemund, P. W. Folding DNA to create nanoscale shapes and patterns. Nature 440, 297–302 (2006).

[3] Zhan, P. et al. Recent advances in DNA origami-engineered nanomaterials and applications. Chemical Reviews 123, 3976–4050 (2023).

[4] Jahnke, K. et al. DNA origami signaling units transduce chemical and mechanical signals in synthetic cells. Advanced Functional Materials 34, 2301176 (2024).

[5] Um, S. H. et al. Enzyme-catalysed assembly of DNA hydrogel. Nature Materials 5, 797–801 (2006).

[6] Takinoue, M. DNA droplets for intelligent and dynamical artificial cells: From the viewpoint of computation and non-equilibrium systems. Interface Focus 13, 20230021 (2023).

[7] Udono, H., Gong, J., Sato, Y. & Takinoue, M. DNA droplets: intelligent, dynamic fluid. Advanced Biology 7, 2200180 (2023).

[8] Gong, J., Tsumura, N., Sato, Y. & Takinoue, M. Computational DNA droplets recognizing miRNA sequence inputs based on liquid–liquid phase separation. Advanced Functional Materials 32, 2202322 (2022).

[9] Tschurikow, X. et al. Amphiphiles formed from synthetic DNA-nanomotifs mimic the stepwise dispersal of transcriptional clusters in the cell nucleus. Nano Letters 23, 7815–7824 (2023).

[10] Hilbert, L. et al. Chromatin-associated condensates as an inspiration for the system architecture of future DNA computers. Annals of the New York Academy of Sciences 1552, 12–28 (2025).

[11] Can, A. E., Ali, A. W., Oelschlaeger, C., Willenbacher, N. & Stoev, I. D. Mechanically tunable DNA hydrogels as prospective biosensing modules. Macromolecular Rapid Communications 2500149 (2025).

[12] Cangialosi, A. et al. DNA sequence–directed shape change of photopatterned hydrogels via high-degree swelling. Science 357, 1126–1130 (2017).

[13] Peng, Y.-H. et al. Dynamic matrices with DNA-encoded viscoelasticity for cell and organoid culture. Nature Nanotechnology 18, 1463–1473 (2023).

[14] Wang, D., Hu, Y., Liu, P. & Luo, D. Bioresponsive DNA hydrogels: beyond the conventional stimuli responsiveness. Accounts of Chemical Research 50, 733–739 (2017).

[15] Gačanin, J., Synatschke, C. V. & Weil, T. Biomedical applications of DNA-based hydrogels. Advanced Functional Materials 30, 1906253 (2020).

[16] Li, F., Lyu, D., Liu, S. & Guo, W. DNA hydrogels and microgels for biosensing and biomedical applications. Advanced Materials 32, 1806538 (2020).

[17] Biffi, S. et al. Phase behavior and critical activated dynamics of limited-valence DNA nanostars. Proceedings of the National Academy of Sciences 110, 15633–15637 (2013).

[18] Brady, R. A., Brooks, N. J., Cicuta, P. & Di Michele, L. Crystallization of amphiphilic DNA C-stars. Nano Letters 17, 3276–3281 (2017).

[19] Saccá, B. & Niemeyer, C. M. Functionalization of DNA nanostructures with proteins. Chemical Society Reviews 40, 5910–5921 (2011).

[20] Sato, Y., Sakamoto, T. & Takinoue, M. Sequence-based engineering of dynamic functions of micrometer-sized DNA droplets. Science Advances 6, eaba3471 (2020).

[21] Zhao, Q.-H., Cao, F.-H., Luo, Z.-H., Huck, W. T. & Deng, N.-N. Photoswitchable molecular communication between programmable DNA-based artificial membraneless organelles. Angewandte Chemie 134, e202117500 (2022).

[22] Leathers, A. et al. Reaction–diffusion patterning of DNA-based artificial cells. Journal of the American Chemical Society 144, 17468–17476 (2022).

[23] Tanase, D. A. et al. Internal phase separation in synthetic DNA condensates. Advanced Science 12, e06275 (2025).

[24] Li, Y., Chen, R., Zhou, B., Dong, Y. & Liu, D. Rational design of DNA hydrogels based on molecular dynamics of polymers. Advanced Materials 36, 2307129 (2024).

[25] Gadzekpo, A., Oprzeska-Zingrebe, E. A., Kozlowska, M., Hilbert, L. & Stoev, I. D. Integrative approaches for DNA sequence-controlled functional materials. Advanced Functional Materials e19573 (2025).

[26] Dans, P. D., Walther, J., Gómez, H. & Orozco, M. Multiscale simulation of DNA. Current Opinion in Structural Biology 37, 29–45 (2016).

[27] Stoev, I. D. et al. On the role of flexibility in linker-mediated DNA hydrogels. Soft Matter 16, 990–1001 (2020).

[28] Weeratunge, H. et al. Bayesian coarsening: rapid tuning of polymer model parameters. Rheologica Acta 62, 477–490 (2023).

[29] Ray, P. et al. Refining coarse-grained molecular topologies: a Bayesian optimization approach. npj Computational Materials 11, 234 (2025).

[30] Ouldridge, T. E., Louis, A. A. & Doye, J. P. Structural, mechanical, and thermodynamic properties of a coarse-grained DNA model. The Journal of Chemical Physics 134, 085101 (2011).

[31] Snodin, B. E. et al. Introducing improved structural properties and salt dependence into a coarse-grained model of DNA. The Journal of Chemical Physics 142, 234901 (2015).

[32] Poppleton, E. et al. oxDNA: coarse-grained simulations of nucleic acids made simple. Journal of Open Source Software 8, 4693 (2023).

[33] Rovigatti, L., Šulc, P., Reguly, I. Z. & Romano, F. A comparison between parallelization approaches in molecular dynamics simulations on GPUs. Journal of Computational Chemistry 36, 1–8 (2015).

[34] Poppleton, E., Romero, R., Mallya, A., Rovigatti, L. & Šulc, P. OxDNA.org: a public webserver for coarse-grained simulations of DNA and RNA nanostructures. Nucleic Acids Research 49, W491–W498 (2021).

[35] Zadeh, J. N. et al. NUPACK: Analysis and design of nucleic acid systems. Journal of Computational Chemistry 32, 170–173 (2011).

[36] Fornace, M. E., Porubsky, N. J. & Pierce, N. A. A unified dynamic programming framework for the analysis of interacting nucleic acid strands: enhanced models, scalability, and speed. ACS Synthetic Biology 9, 2665–2678 (2020).

[37] Dirks, R. M., Bois, J. S., Schaeffer, J. M., Winfree, E. & Pierce, N. A. Thermodynamic analysis of interacting nucleic acid strands. SIAM Review 49, 65–88 (2007).

[38] Hoffmann, M., Fröhner, C. & Noé, F. ReaDDy 2: Fast and flexible software framework for interacting-particle reaction dynamics. PLoS Computational Biology 15, e1006830 (2019).

[39] Conrad, N., Kennedy, T., Fygenson, D. K. & Saleh, O. A. Increasing valence pushes DNA nanostar networks to the isostatic point. Proceedings of the National Academy of Sciences 116, 7238–7243 (2019).

[40] Palombo, G., Weir, S., Michieletto, D. & Gutiérrez Fosado, Y. A. Topological linking determines elasticity in limited valence networks. Nature Materials 24, 454–461 (2025).

[41] Xing, Z. et al. Microrheology of DNA hydrogels. Proceedings of the National Academy of Sciences 115, 8137–8142 (2018).

[42] Fernandez-Castanon, J., Bianchi, S., Saglimbeni, F., Di Leonardo, R. & Sciortino, F. Microrheology of DNA hydrogel gelling and melting on cooling. Soft Matter 14, 6431–6438 (2018).

[43] Cohen, S. R., Banerjee, P. R. & Pappu, R. V. Direct computations of viscoelastic moduli of biomolecular condensates. The Journal of Chemical Physics 161, 0951030 (2024).

[44] Xing, Z., Ness, C., Frenkel, D. & Eiser, E. Structural and linear elastic properties of DNA hydrogels by coarse-grained simulation. Macromolecules 52, 504– 512 (2019).

[45] Fosado, Y. A. G. Nanostars planarity modulates the rheology of DNA hydrogels. Soft Matter 19, 4820–4828 (2023).

[46] Ajiyel, H. et al. Mechanical response of a simple DNA nanostar hydrogel: symptoms of disorder and glassy emergence of solidity. arXiv preprint 2603.20119 (2026).

[47] Song, J., Holten-Andersen, N. & McKinley, G. H. Non-Maxwellian viscoelastic stress relaxations in soft matter. Soft Matter 19, 7885–7906 (2023).

[48] Gomez, S. S. & Rovigatti, L. Diffusion, viscosity, and linear rheology of valence-limited disordered fluids. The Journal of Chemical Physics 160, 184901 (2024).

[49] Aime, S., Cipelletti, L. & Ramos, L. Power law viscoelasticity of a fractal colloidal gel. Journal of Rheology 62, 1429–1441 (2018).

[50] Hu, Y. et al. Quantifying t cell receptor mechanics at membrane junctions using DNA origami tension sensors. Nature Nanotechnology 19, 1674–1685 (2024).

[51] Bukina, V. & Božič, A. Context-dependent structure formation of hairpin motifs in bacteriophage MS2 genomic RNA. Biophysical Journal 123, 3397–3407 (2024).

[52] Yang, L., Pecastaings, G., Drummond, C. & Elezgaray, J. Driving DNA nanopore membrane insertion through dipolar coupling. Nano Letters 24, 13481–13486 (2024).

[53] Liu, X. et al. A lumen-tunable triangular DNA nanopore for molecular sensing and cross-membrane transport. Nature Communications 15, 7210 (2024).

[54] Mogheiseh, M. & Hasanzadeh Ghasemi, R. Design and simulation of a wireframe DNA origami nanoactuator. The Journal of Chemical Physics 161, 045101 (2024).

[55] Walker-Gibbons, R., Zhu, X., Behjatian, A., Bennett, T. J. & Krishnan, M. Sensing the structural and conformational properties of single-stranded nucleic acids using electrometry and molecular simulations. Scientific Reports 14, 20582 (2024).

[56] Kaufhold, W. oxDNA tutorial (2020). URL https://github.com/WillTKaufhold1/oxdna-tutorial.

[57] Ouldridge, T. E., Louis, A. A. & Doye, J. P. Extracting bulk properties of self-assembling systems from small simulations. Journal of Physics: Condensed Matter 22, 104102 (2010).

[58] Wu, Y.-Y., Bao, L., Zhang, X. & Tan, Z.-J. Flexibility of short DNA helices with finite-length effect: from base pairs to tens of base pairs. The Journal of Chemical Physics 142, 125103 (2015).

[59] Geggier, S., Kotlyar, A. & Vologodskii, A. Temperature dependence of DNA persistence length. Nucleic Acids Research 39, 1419–1426 (2011).

[60] Theodorakopoulos, N. & Peyrard, M. Base pair openings and temperature dependence of DNA flexibility. Physical Review Letters 108, 078104 (2012).

[61] Balandat, M. et al. BoTorch: A Framework for Efficient Monte-Carlo Bayesian Optimization. In Advances in Neural Information Processing Systems 33 (2020).

[62] Paszke, A. et al. Pytorch: An imperative style, highperformance deep learning library. In Advances in Neural Information Processing Systems 32, 8024– 8035 (Curran Associates, Inc., 2019).

[63] Frazier, P. I. A tutorial on bayesian optimization. arXiv preprint 1807.02811 (2018).

[64] Wolfe, B. R., Porubsky, N. J., Zadeh, J. N., Dirks, R. M. & Pierce, N. A. Constrained multistate sequence design for nucleic acid reaction pathway engineering. Journal of the American Chemical Society 139, 3134–3144 (2017).

[65] Wolfe, B. R. & Pierce, N. A. Sequence design for a test tube of interacting nucleic acid strands. ACS Synthetic Biology 4, 1086–1100 (2015).

[66] Webber, M. J. & Tibbitt, M. W. Dynamic and reconfigurable materials from reversible network interactions. Nature Reviews Materials 7, 541–556 (2022).

[67] Rouse Jr, P. E. A theory of the linear viscoelastic properties of dilute solutions of coiling polymers. The Journal of Chemical Physics 21, 1272–1280 (1953).

[68] Zimm, B. H. Dynamics of polymer molecules in dilute solution: viscoelasticity, flow birefringence and dielectric loss. The Journal of Chemical Physics 24, 269–278 (1956).

[69] Doi, M., Edwards, S. F. & Edwards, S. F. The theory of polymer dynamics, vol. 73 (Oxford University Press, 1988).

[70] Wong, C. P. J. & Choi, P. Numerical calculation of intrinsic viscosity of star and ring polymers using a modified Zimm model. The Journal of Chemical Physics 163, 154901 (2025).

[71] Jurjiu, A. & Galiceanu, M. Dynamics of a polymer network modeled by a fractal cactus. Polymers 10, 787 (2018).

[72] Ferry, J. D. Viscoelastic properties of polymers (John Wiley & Sons, Inc., 1980).

[73] Alshareedah, I. et al. Sequence-specific interactions determine viscoelasticity and ageing dynamics of protein condensates. Nature Physics 20, 1482–1491 (2024).

[74] Rovigatti, L., Bomboi, F. & Sciortino, F. Accurate phase diagram of tetravalent DNA nanostars. The Journal of Chemical Physics 140, 154903 (2014).

[75] Dupuis, N. F., Holmstrom, E. D. & Nesbitt, D. J. Single-molecule kinetics reveal cation-promoted DNA duplex formation through ordering of single-stranded helices. Biophysical Journal 105, 756–766 (2013).

[76] Flory, P. J. Molecular size distribution in three dimensional polymers. i. gelation1. Journal of the American Chemical Society 63, 3083–3090 (1941).

[77] Stockmayer, W. H. Theory of molecular size distribution and gel formation in branched polymers II. General cross linking. The Journal of Chemical Physics 12, 125–131 (1944).

[78] Ewoldt, R. H., Johnston, M. T. & Caretta, L. M. Experimental Challenges of Shear Rheology: How to Avoid Bad Data, 207–241 (Springer New York, New York, NY, 2015).

[79] Winter, H. H. & Chambon, F. Analysis of linear viscoelasticity of a crosslinking polymer at the gel point. Journal of Rheology 30, 367–382 (1986).

[80] Winter, H. H. Gel Point, 1–15 (John Wiley & Sons, Ltd, 2016).

[81] Mohar, B., Alavi, Y., Chartrand, G. & Oellermann, O. The Laplacian spectrum of graphs. Graph Theory, Combinatorics, and Applications 2, 12 (1991).

[82] Zhang, X.-D. The Laplacian eigenvalues of graphs: a survey. arXiv preprint 1111.2897 (2011).

[83] Zhang, R., Mitra, G., Ghosh, S. & Pappu, R. V. Computational rheometry for modeling viscoelasticity and mechanical responses of biomolecular condensates. Biophysical Journal 125, 1979–1995 (2026).

[84] Lewis, C. L., Stewart, K. & Anthamatten, M. The influence of hydrogen bonding side-groups on viscoelastic behavior of linear and network polymers. Macromolecules 47, 729–740 (2014).

[85] Ginzburg, V. V., Chatterjee, T., Nakatani, A. I. & Van Dyk, A. K. Oscillatory and steady shear rheology of model hydrophobically modified ethoxylated urethane-thickened waterborne paints. Langmuir 34, 10993–11002 (2018).

[86] Le Roy, H. et al. Valence can control the nonexponential viscoelastic relaxation of multivalent reversible gels. Science Advances 10, eadl5056 (2024).

[87] Jiang, S. & Webb, M. A. Generative active learning across polymer architectures and solvophobicities for targeted rheological behavior. npj Computational Materials (2025).

[88] Ding, L.Carrillo, J.-M. & Do, C. ToPolyAgent: AI agents for coarse-grained bead-spring topological polymer simulations. Digital Discovery 5, 901–909 (2026).

[89] Jiang, S., Dieng, A. B. & Webb, M. A. Property-guided generation of complex polymer topologies using variational autoencoders. npj Computational Materials 10, 139 (2024).

[90] Nehil-Puleo, K. & Yang, Z. J. FiberForge: enabling high-throughput simulations of the mechanical properties of helical fibrils. Digital Discovery (2026).

[91] Mao, J., Xi, Y., Zadeh, A. S., Liu, A. P. & Ferguson, A. L. Computational design of polypeptide-based compartments for synthetic cells. Digital Discovery (2026).

[92] Walczak, M. et al. Responsive core-shell DNA particles trigger lipid-membrane disruption and bacteria entrapment. Nature Communications 12, 4743 (2021).

[93] Kengmana, E., Ornelas-Gatdula, E.Chen, K.-L. & Schulman, R. Spatial control over reactions via localized transcription within membraneless DNA nanostar droplets. Journal of the American Chemical Society 146, 32942–32952 (2024).

[94] Stoev, I. D., Caciagli, A., Mukhopadhyay, A., Ness, C. & Eiser, E. Bulk rheology of sticky DNA-functionalized emulsions. Physical Review E 104, 054602 (2021).

[95] Cao, D., Xie, Y. & Song, J. DNA hydrogels in the perspective of mechanical properties. Macromolecular Rapid Communications 43, 2200281 (2022).

