## Supplementary Information for "Bayesian optimisation and graph-based rheology enable sequence-dependent modelling of DNA materials"

#### S1 oxDNA simulations

##### S1.1 Simulation details

Conformational statistics for a single nanomotif were gathered by running molecular-dynamics (MD) simulations in a periodic box of 50x50x50nm. After $1.1 \cdot 10^6$ equilibration steps, $5 \cdot 10^7$ production steps were performed in the NVT ensemble at 20°C and 0.15M NaCl. All other parameters followed established literature standards. [1, 2]

To assess the inter-arm flexibility of two hybridised nanomotif arms, we fused two nanomotifs through their sticky ends. For this purpose, an additional potential between the sticky ends was introduced to accelerate hybridisation. Two previously equilibrated nanomotifs were placed in a box (100x100x100nm) and fused over $10^7$ MD steps. Subsequently, the auxiliary potential was removed and the system was simulated for another $5 \cdot 10^7$ steps under the same conditions as the single-nanomotif run.

##### S1.2 Angle evaluation of oxDNA trajectories

To quantify nanomotif flexibility, groups of typically ten base pairs ($\approx$ one helical turn) were used to define coordinate centres, effectively treating them as coarse-grained beads. Vectors were constructed between these centres and employed to calculate angles within and between motif arms. The positions of nucleotide groups were extracted from oxDNA trajectories using their unique nucleotide IDs, and centres of mass were computed from these positions. We tested several coarse-graining schemes and observed that the resulting angles were consistent, provided that each group contained enough nucleotides to align the vectors along the DNA helix. The grouping strategy is illustrated in Figure S2A. Angle histograms were generated from the final 47,000 configurations sampled during the production MD run. Energy profiles remained stable throughout, confirming that the simulations were equilibrated (Figure S1A,B).

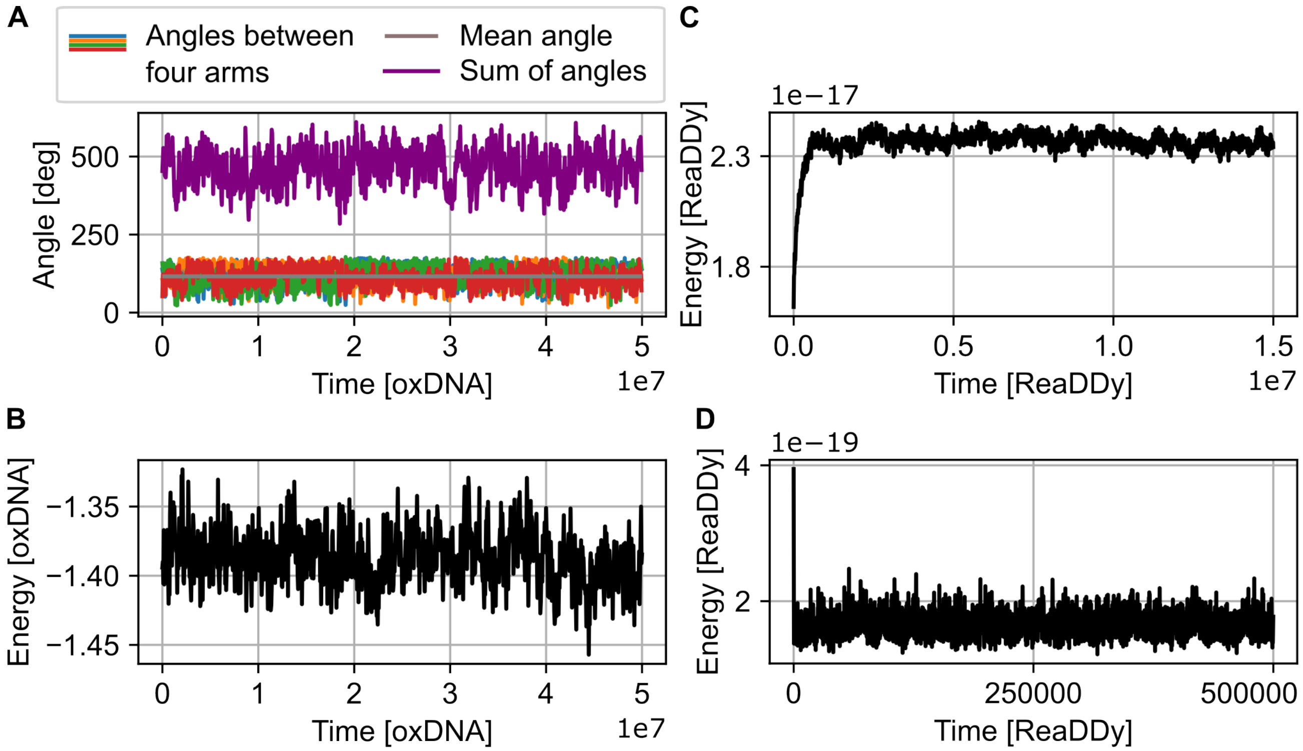


Figure S1: Energies and angles after equilibration. (A) Angles between the four nanomotif arms as well as (B) energies during the production run. Every tenth step shown. (C) Example of energy as a function of time during network assembly of 100 nanomotifs implemented in ReaDDy. Equilibration phase not shown. (D) Energies during example ReaDDy simulation to sample conformations of a single nanomotif. The first 300 steps with a strong drop in energy due to the initial relaxation after placing the motif in the box were excluded.

##### S1.3 Sampling of sticky end hybridisation

Umbrella sampling was used to evaluate the thermodynamics of sticky end hybridisation. We investigated hairpin formation within one sticky end and hybridisation between two sticky ends.

Sampling of hybridisation events used $10^5$ equilibration and $10^7$ production steps, where dozens of transitions between fully open and fully hybridised configurations were observed. Two improve statistics, 6 independent repeats were carried out. States were defined by proximity between base pairs and presence of hydrogen bonds. To sample each state sufficiently, weights for umbrella sampling were iteratively updated by inverting the observed frequency. The code to derive weights and produce free energy curves is based on a tutorial by Will Kaufhold [3] available on GitHub (`https://github.com/WillTKaufhold1/oxdna-tutorial`).

For sampling of hairpin formation, we noticed that inclusion of an arm segment yielded statistics matching those from the full MD run that we used to sample conformations (Figure S3). This indicated that the configurational constrain due to the presence of an arm fragment influences the free energy. We therefore also included the arm fragment when sampling hybridisation between two sticky ends.

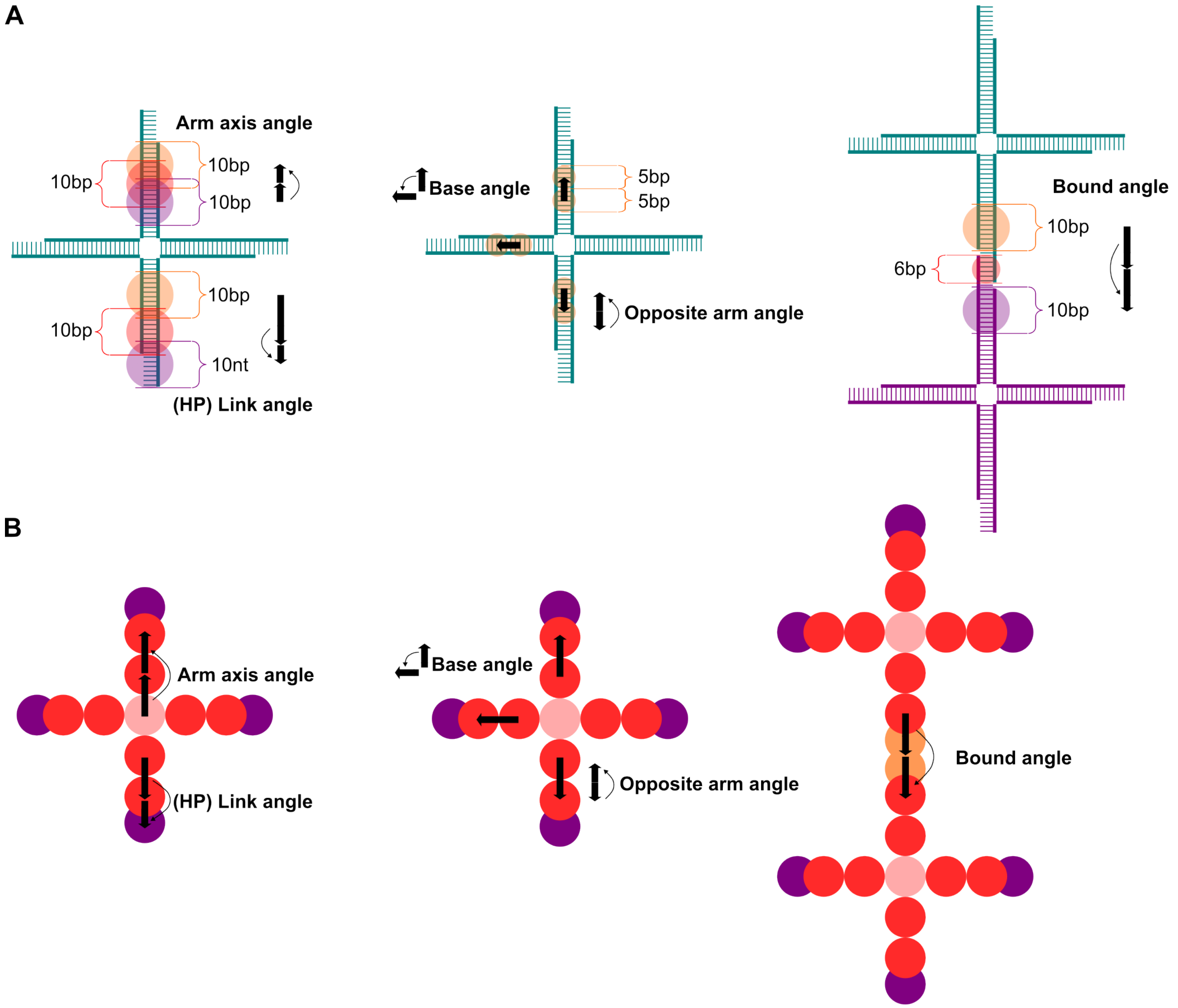


Figure S2: Angle definitions in oxDNA and ReaDDy (A) Groups of nucleotides were used to define centre coordinates for vectors. Nucleotides close to the centre were avoided as the DNA structure is highly flexible in this region. When possible, groups consisted of 10 base pairs to align the vectors close to the helix axis. For base and opposite arm angles, vectors were aligned by spitting 10 base pairs in the central arm part to define base and tip of the vector. Modifying the definitions for the groups did not strongly affected resulting angle distributions, as long as physically meaningful parts of the motif were grouped. (B) Angle definitions in the bead-spring model followed the oxDNA definitions but used the 13 beads as reference particles. The final beads (purple) representing the sticky end were shifted inwards, so that fusion of two beads created a distance corresponding to 8 base pairs, consistent with the hybridisation of two sticky ends.

### S2 ReaDDy simulations

#### S2.1 Simulation details and parameter space for optimisation

Nanomotifs were coarse-grained into 13 beads. The diffusion constant for beads in the bead-spring model was estimated based on measurements of DNA oligo mobility in solution [4, 5] and was comparable to prediction from the Einstein-Smoluchowski [6, 7] equation for a sphere ($\approx 0.101$ versus $0.158\text{nm}^2/\text{ns}$). The spacing between beads roughly corresponded to 8 base pairs ($\approx 2.7\text{nm}$). Electrostatic repulsion between beads was modelled with Debye-Hueckel potentials and used the

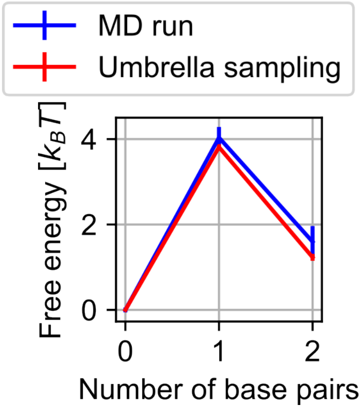


Figure S3: Free energy of hairpin formation. Both umbrella sampling and statistics from the MD run produced profiles within the standard error of each other.

| Parameter | Range of values | Optimised value | Unit |
|---|---|---|---|
| $k_{arm}$ | $[10^{-24}, 10^{-18}]$ | $2.033 \cdot 10^{-19}$ | J/rad$^2$ |
| $k_{base}$ | $[10^{-24}, 10^{-18}]$ | $1 \cdot 10^{-24}$ | J/rad$^2$ |
| $\Theta_{eq\_base}$ | [75,120] | 120 | deg |
| $k_{oppa}$ | $[10^{-24}, 10^{-18}]$ | $5.836 \cdot 10^{-21}$ | J/rad$^2$ |
| $\Theta_{eq\_oppa}$ | [135,180] | 164.497 | deg |
| $k_{link}$ | $[10^{-24}, 10^{-18}]$ | $4.231 \cdot 10^{-20}$ | J/rad$^2$ |
| $k_{HP}$ | $[10^{-24}, 10^{-18}]$ | $5.489 \cdot 10^{-20}$ | J/rad$^2$ |
| $k_{bound}$ | $[10^{-24}, 10^{-18}]$ | $1.572 \cdot 10^{-20}$ | J/rad$^2$ |

Table S1: Parameter ranges for tunable force field parameters in bead-spring model

same effective charge per nucleotide ($q_e$=0.815) used in the oxDNA model.[8] The Debye-Hückel potentials used a distance cut-off set to a value of 4 times the Debye screening length ensuring that potential values had dropped sufficiently before the cut-off. Beads were connected with harmonic potentials, with spring constants set to $2 \cdot 10^{-19}$J/nm$^2$ and $10^{-19}$J/nm$^2$ for double and single stranded DNA, in line with values used in previous simulations and found in experimental studies. [9]

Harmonic spring potentials $V_{angle} = k_{angle}(\Theta - \Theta_0)^2$ between three beads were used as tunable force field, capable of producing a wide range of motif conformational statistics. Table S1 contains the parameter ranges we investigated with ReaDDy simulations. We chose wide ranges for spring constants and equilibrium angles to screen a large space of potential behaviour. The lowest spring constants were effectively zero while the highest spring constants resulted in rigid structures. Both rigid and extremely flexible nanomotifs were strongly inhibited in their capability to form networks, either through steric constrains or by collapsing of few motifs into self-hybridised "blobs". To obtain angle distributions, nanomotifs were placed in a large simulation box (150x150x150nm) and simulated for $5 \cdot 10^5$ time steps using overdamped Langevin dynamics (Brownian dynamics). We used a time step of $10^{-2}$ ns and the first $3 \cdot 10^4$ steps as equilibration phase, yielding converging angle distributions and stable energies (Figure S1D). Angles were defined between vectors aligned to the coarse-grained beads (Figure S2B). Bead positions were retrieved via particle IDs. For each angle a distribution was generated and compared to the corresponding coarse-grained angle distribution in the oxDNA model. The optimised parameters found by the methods described in the main text can be found in Table S1. Note that optimisation set the base angle effectively to zero, indicating that the opposite arm angle was sufficient to reproduce oxDNA reference angle distributions.

##### S2.2 Simulating the assembly of nanomotif networks

Networks between 100 nanomotifs were assembled via MD simulations in a box with 69.7x69.7x69.7nm, matching experimental concentrations.[10] Network formation happened by introducing a reaction that fused sticky ends with a certain probability, once they were close together ($\leq$2.7nm) in space. Additionally, both sticky ends had to be in an open and not a hairpin configuration. Ratios of forward and backward reaction steps were used to simulate sequence and condition dependent differences in network connectivity. Internally, reactions were chosen with kinetic Monte-Carlo sampling, with a probability $p_{reaction} = 1 - e^{-k \cdot \Delta t}$ for a reaction with rate $k$ to occur given an integration time step $\Delta t$. [11] All sticky ends could be involved in reactions independently from each other and reactions were evaluated every time step. Hairpin formation can happen irrespective of the presence of other nanomotifs and was frequently observed during MD simulations with oxDNA. We chose a forward rate for hairpin formation of $r_{HP} \cdot \Delta t = 5 \cdot 10^{-5}$. The backward reaction rate $r_{Link}$ was chosen to match the equilibrium fraction of time spent in hairpin configuration, according to oxDNA (see main text). With this choice, hairpins were formed and opened at a faster time scale than nanomotifs would bind and unbind, in line with more stable hybridisation between two sticky ends. With a fusing rate between two sticky ends of $r_{Fuse} \cdot \Delta t = 5 \cdot 10^{-4} nm^{-3}$, the nanomotif network was gradually assembled during simulations. Different splitting rates $r_{Split}$ (see main text) were used to simulate differences in network connectivity. For non-zero splitting rates, nanomotifs could become unbound and diffuse to a different location in the simulation box, indicative of liquid-like internal reorganisation experimentally observed for nanomotifs.[12] We tested different absolute values for the rates but found that this had little effect on the observed network properties, indicating that they are mostly determined by equilibrium properties. For simulations that were evaluated for inverse design, no hairpin formation was simulated, as the sequence is not known a priori. The simulated rates can thus be interpreted as effective binding and unbinding rates that implicitly include hairpin formation.

##### S2.3 Graph interpretation of ReaDDy simulations

To apply graph-based rheology methods to our nanomotif simulations, we extracted graph Laplacians from simulations of networks of nanomotifs. As discussed in the main text, hybridisation between nanomotifs was modelled with a fusion reaction. ReaDDy treats resulting networks of nanomotifs as topologies with nodes (particles) and edges (bonds between particles). From the record of topologies, a graph Laplacian can be generated for each time step. First, the centre particle of each nanomotif was selected. From this centre particle, 4 arms with potentially fused sticky ends emanate. For each of the 4 sticky ends of nanomotif $i$, we checked whether a bond with any sticky ends of other nanomotifs was present. If a bond with a sticky end of nanomotif $j$ was found, this yielded an entry of $L(i, j) = -1$ in the graph Laplacian $L$. The eigenvalues of this matrix were used to calculate G' and G" as discussed in the main text and the literature [13]. As described in the main text, multiple time steps and corresponding G' and G" curves were averaged from multiple repeats.

#### S3 Extended results of Bayesian force field optimisation

To validate the Bayesian optimisation strategy and the choice of hyperparameters, we selected 13 random parameter combinations for the coarse-grained bead-spring model and used their angle distributions as validation targets (Figure S4A). Due to the correlations between free parameters (see main text), we compared split optimisation (optimise angles within arms first, then between arms) and combined optimisation (optimise angles within and between arms simultaneously). Sticky end

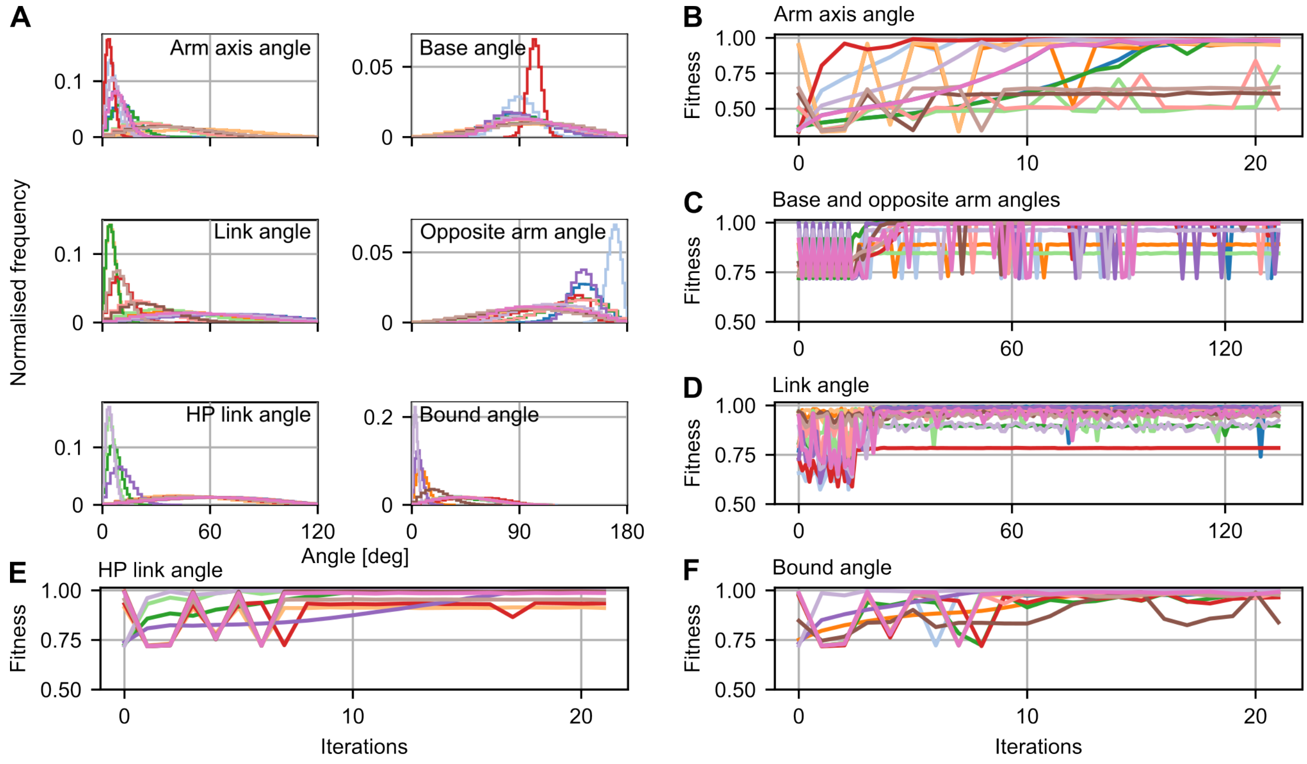


Figure S4: Target angle distributions and fitness during validation. (A) 13 validation targets were obtained from random parameter combinations using the bead-spring model. Fitness versus iterations shown for split optimisation of the arm axis angle (B), the coupled base and opposite arm angle (C), the link angle (D), the link angle in hairpin configuration (E), and the bound arm angle (F). All data was averaged from five simulations per parameter set.

angles and angles between fused arms were optimised separately as they did not correlate strongly with other angles. 1D problems used two initial trial simulations followed by 20-30 optimisation iterations. The 4D problem of the split optimisation used 16+120 initial and optimisation iterations, the 5D problem of the combined optimisation used 32+120 initial and optimisation iterations. The number of initial simulations allowed to probe the extreme "corners" of the search space before starting the optimisation. With the selected number of iterations, the optimisation converged for the 13 validation cases, usually to a fitness approaching 1 (Figure S4B-F). We quantified the performance of both optimisation strategies during validation by computing the Wasserstein distance (Figure S5C) and conducting a Kolmogorov–Smirnov test (KS) test between optimised and target angle distributions (Figure S5B).

Figure S5A shows an example of successful optimisation of the arms axis angle, where the Wasserstein distance between target and optimised angle distribution was $\approx 0.01°$ for both optimisation strategies. In this example, the link angle was only successfully optimised with the split optimisation strategy (Wasserstein distance $\approx 0.01°$) whereas a notable deviation from the target (Wasserstein distance $\approx 0.26°$) persisted after combined optimisation.

In general, the optimisation performed best on the base angle, where none of the optimised angle distributions were significantly different from their targets, irrespective of the optimisation strategy. The two strategies performed well on all other angle distributions with similar fractions (77%-100%) of optimised angle distributions not significantly different from their targets (Figure S5B). A histogram of the Wasserstein distances from all 13 optimisation targets for both methods can be found in Figure S5C, again showing that the two strategies performed similarly.

Moving to oxDNA target angle distributions, Wasserstein distances between optimised angle distributions and oxDNA targets revealed that the combined optimisation performed worse than

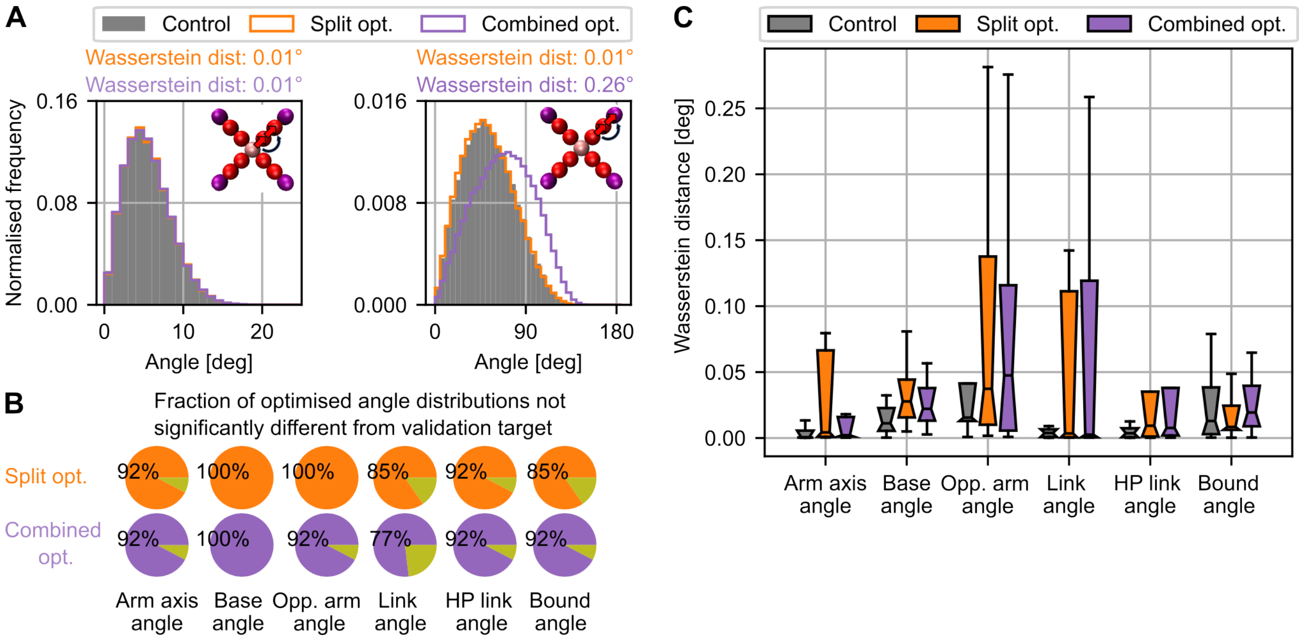


Figure S5: Validating Bayesian optimisation. (A) Examples of successful optimisation of the arm axis angle (left histogram) and unsuccessful optimisation when using the combined optimisation strategy for the link angle (right histogram). (B) 13 random parameter combinations were used as validation targets. A Kolmogorov–Smirnov test (KS) test between optimised and target angle distributions was used to quantify what fraction of angle distributions was not significantly different from their target. (C) Boxplots of Wasserstein distances between optimised and target angle distributions from 13 validation targets. Notched regions are 95% bootstrap confidence intervals. Control histograms were obtained with optimal parameters, all angle distributions were averaged from five simulations.

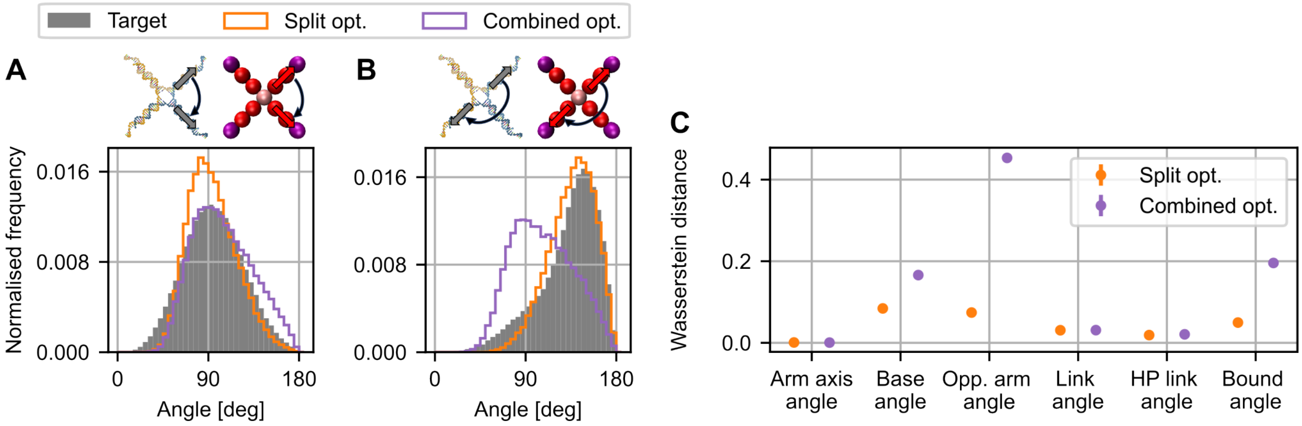


Figure S6: Comparing optimisation strategies. Optimised base angle (A) and opposite arm angle (B) distributions from both strategies. (C) Wasserstein distances between optimised and oxDNA target angle distributions. Optimised angle distributions averaged from five repeats. Mean and standard error of the mean in (C) obtained from three repeats each averaging five simulations.

split optimisation on the base, opposite and bound angle, but almost identical for all other angles (Figure S6). The split optimisation produced a close match with Wasserstein distances below $\approx 0.1°$. Notable deviations only persisted for the base and opposite angles, likely because the bead-spring force field cannot capture the full complexity of the central nanomotif structure. Figure S7 displays the fitness convergence for the 4D split optimisation using one Gaussian process. $k_{link}$ of the sticky end was optimised with a second Gaussian process applied to the same set of trial simulations.

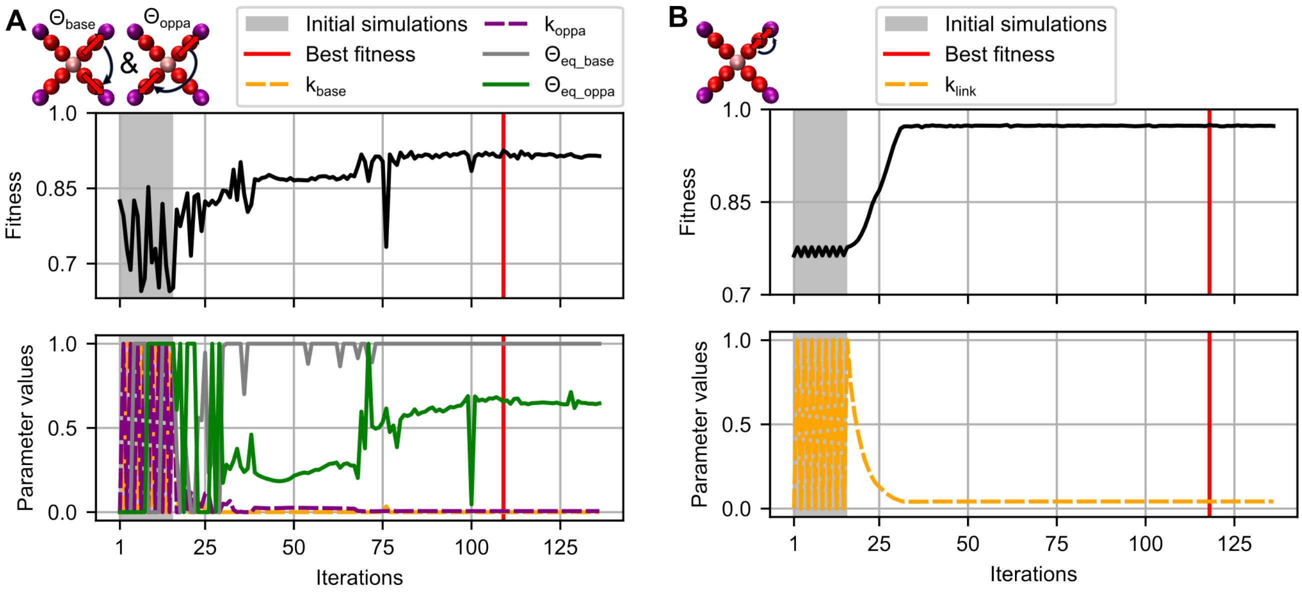


Figure S7: Optimising fitness by tuning bead-spring model parameters. (A) In the split optimisation approach, the four parameters $k_{base}$, $k_{oppa}$, $\theta_{eq\_base}$ and $\theta_{eq\_oppa}$ were optimised simultaneously. After 108 iterations, the best fitness for the base and opposite arm angle was found (red line). (B) A second Gaussian process using the same trial simulations was used to optimise $k_{link}$. The fitness of the link angle converged after $\approx 40$ iterations, however, the best fitness was found in iteration 117, as angle distributions and resulting fitness scores fluctuated. Simulations used to initialise the Gaussian process model were carried out in the grey area. Data from five simulations was averaged for each set of parameters.

#### S4 Sticky end designs for evaluated fractions of bound sticky end

We applied Bayesian optimisation and NUPACK to find potentials DNA sequences, salt and temperature conditions for the lowered fraction of bound sticky ends ($\approx 61\%$) we evaluated in the main text. These serve to illustrate that minor modifications of the DNA system can tune the expected fraction of bound sticky ends. As discussed in the main text, NUPACK predictions do not consider the fact that sticky ends are bound to nanomotif arms. Consequently, actual fractions of bound sticky ends might be shifted. In future work, this potential deviation could be accounted for with an additive constant to the free energies, which could be obtained by comparing NUPACK results to oxDNA predictions and experiments.

| Sequence | Temp. [°C] | [NaCl][M] |
|---|---|---|
| ACTAGT | 28.8 | 0.192 |
| GAATTC | 20.6 | 0.131 |
| TGTACA | 29.7 | 0.127 |

Table S2: Sticky end sequences, salt and temperature conditions for a bound fraction of $\approx 0.61$. The concentration of bivalent ions was set to zero.

#### S5 Generation of theoretical graphs and quantification of predictions for n-armed nanomotifs

To generate random graphs of networked materials we first deterministically constructed a simple graph with N nodes and exactly m undirected edges per node. This graph corresponded to the maximum possible connectivity in the network. From this graph, we randomly selected a subset of

3-armed motif

| | Graph-based | | Conventional | |
|---|---|---|---|---|
| | $R^2$ | MAE, (log) | $R^2$ | MAE, (log) |
| G' | 0.990 (0.908) | 0.108 (0.375) | 0.953 | 0.135 |
| G" | 0.915 (0.839) | 0.720 (0.632) | 0.833 | 0.924 |

3-armed motif, microrheology

| | Graph-based | | Conventional | |
|---|---|---|---|---|
| | $R^2$ | MAE, (log) | $R^2$ | MAE, (log) |
| G' | 0.997 (0.963) | 0.053 (0.249) | 0.992 | 0.125 |
| G" | 0.971 (0.818) | 0.046 (0.115) | 0.749 | 0.153 |

4-armed motif

| | Graph-based | | Conventional | |
|---|---|---|---|---|
| | $R^2$ | MAE, (log) | $R^2$ | MAE, (log) |
| G' | 0.986 (0.989) | 0.107 (0.135) | 0.873 | 0.152 |
| G" | 0.943 (0.946) | 0.844 (0.819) | 0.836 | 1.102 |

5-armed motif

| | Graph-based | | Conventional | |
|---|---|---|---|---|
| | $R^2$ | MAE, (log) | $R^2$ | MAE, (log) |
| G' | 0.975 | 0.114 | 0.885 | 0.184 |
| G" | 0.927 | 0.939 | 0.837 | 1.152 |

6-armed motif

| | Graph-based | | Conventional | |
|---|---|---|---|---|
| | $R^2$ | MAE, (log) | $R^2$ | MAE, (log) |
| G' | 0.955 | 0.092 | 0.867 | 0.240 |
| G" | 0.940 | 1.031 | 0.869 | 1.228 |

Table S3: Similarity between rescaled moduli from graph-based or conventional Maxwell model and experimental data. Values in brackets did not use the cutoff relaxation time discussed in the main text. Mean absolute errors (MAE) were integrated in log space.

edges to realise graphs with varying connectivity and randomised the structure further by 5000 2-switches. This step sampled from the space of all possible simple graphs with the specified degree. We found that this strategy produced the same trend of crossover frequencies as a function of added edges that we obtained from the simulations. Alternatively, we tried a method that first generated all possible edges in the network and then sampled a subset of them, which yielded similar G' and G" results, but sometimes failed to achieve full connectivity due to the greedy sampling strategy. Since our simulations rarely contained more than one hybridised arm between two nanomotifs and no permanent hybridisation between two arms of one motif, we excluded parallel edges and self-loops in our theoretical graphs.

We compared predictions for G' and G" from the conventional, single relaxation time Maxwell model as well as predictions from the generalised, graph-based model to experiment data (Table S3). [10, 14] Predictions were rescaled to match the crossover point between G' and G" of the experiment. For networks further away from the isostatic point, similarity to the experimental data was measured with and without cutoff relaxation time, showing how introduction of such a cutoff improved predictions. Generally, the graph-based method using a spectrum of relaxation times from networks with 800 nodes described the experimental data well and provided better predictions than the single relaxation time approach.

#### S6 Hydrodynamic interactions in classical polymer models and simulation results of 3-armed nanomotifs

In the classical Rouse picture, [15] a spectrum of relaxation times $\tau_p \propto p^{-2}$ emerges as sections of a chain of beads connected by springs relax. (Figure S8A) The relaxation times in the Rouse picture, which neglects hydrodynamic interactions, lead to the characteristic scaling of the storage and loss modulus in Figure S8C, with $\omega^{1/2}$ for intermediate frequencies. Hydrodynamic interactions between polymer beads are considered in the Zimm picture (Figure S8B), [16] leading to additional coupling and a different scaling of relaxation times $\tau_p \propto p^{-1.5}$ and a scaling $\omega^{2/3}$ of the moduli. Note that the Zimm model assumes Gaussian chain statistics, which is not imposed in the graph based method discussed in the main text. [17]

Under the assumption of Gaussian chain statistics, the entries of the mobility matrix are given by: [17]

$$H_{nm} = \begin{cases} \dfrac{1}{6\pi\eta a} & \text{if } n = m, \\ \dfrac{1}{6\pi\eta b}\dfrac{1}{\sqrt{6\pi|n-m|}} & \text{if } n \neq m, \end{cases} \tag{1}$$

For figure S8C, we set $\eta = 1$,the Kuhn length $b = 1$ and the bead radius $a = 0.25$.

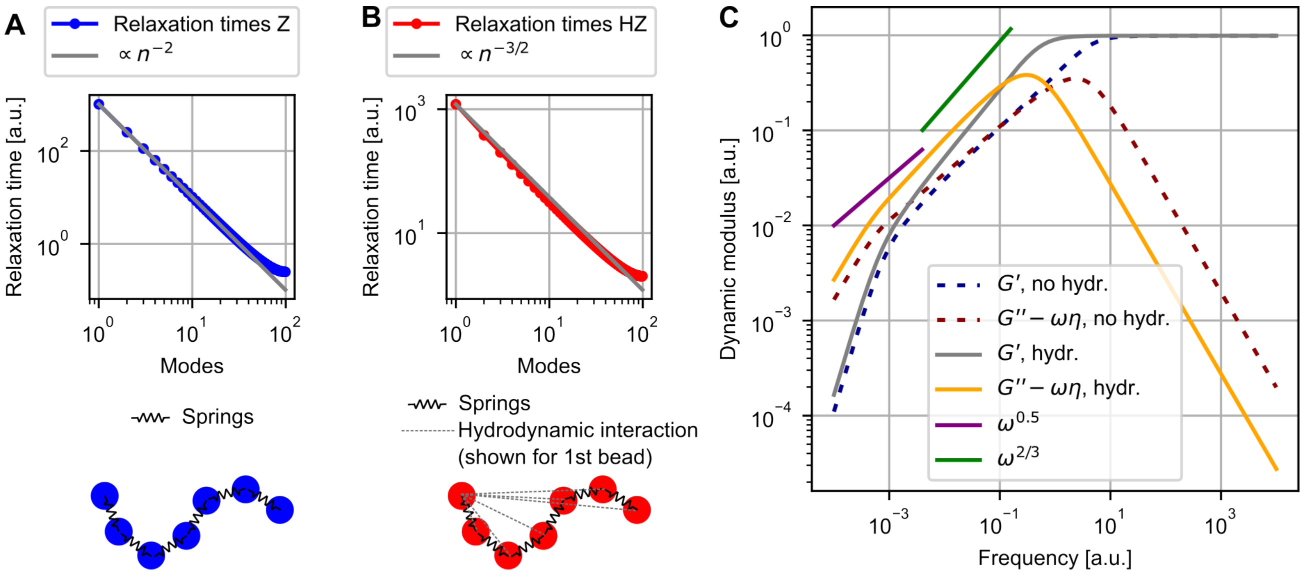


Figure S8: Dynamic moduli predicted by Rouse and Zimm polymer models. (A) Relaxation times for a polymer with 100 beads in the Rouse picture. (B) Relaxation times for a polymer with 100 beads in the Zimm picture. (C) Storage and loss moduli for both Zimm and Rouse polymers.

To assess whether the discrepancy between measured and predicted moduli for three-armed nanostars discussed in the main text arises from the assumption of an ideal, perfectly connected network of degree three, we analysed the graph structure obtained from simulations of 3-armed nanostars. (Figure S9B) These simulations were optimised using the same workflow as in the main text (oxDNA simulations followed by Bayesian parameter tuning to reproduce angle distributions) but used a slightly different sequence design (found in [12]) at 40°C and a salt concentration of 700 mM, and were originally developed for a separate study. To assemble network of 102 motifs in a box (69.7x69.7x69.7nm) with high connectivity, we ran simulations without hairpin formation, and a fusing rate of $r_{Fuse} \cdot \Delta t = 1 \cdot 10^{-3} nm^{-3}$. We ran $2 \cdot 10^6$ equilibration steps, $1.5 \cdot 10^7$ steps with

$r_{split} \cdot \Delta t = 2 \cdot 10^{-8}$ followed by $1.5 \cdot 10^7$ steps with $r_{split} \cdot \Delta t = 0$ resulting in a fraction of bound sticky ends of $0.975 \pm 0.002$. Jupyter notebooks containing all the simulation and parametrisation steps are available on GitHub. To qualitatively probe whether the resulting network graphs differ from the theoretical ones, we adjusted the simulated temperature, salt and nanostar concentrations to the conditions used in the main text and applied the same viscoelastic modulus analysis. To account for hydrodynamic interactions, we assumed an effective radius 3/4 that of the 4-armed nanostar, yielding 3.525 nm, which is smaller than the hydrodynamic radius of 4.5 nm measured in [18]. The smaller radius approximated the experimental reference marginally better. The resulting storage and loss moduli, with and without hydrodynamic interactions are shown in Figure S9A. Compared to predictions from theoretical graphs, graphs extracted from simulations with hydrodynamic interactions approximated the experimental reference slightly better, but still showed persistent deviations. This result suggests that the observed discrepancy between experimental and predicted moduli is unlikely to originate from deviations in the simulated network topology alone. Instead, it may reflect a systematic underestimation of the effective connectivity in the experimental hydrogel network. Such an underestimation could arise from the use of Rouse and Zimm models, which neglect excluded volume effects and topological constraints such topological linking by rings of linked nanostars, which has recently been identified as significant contributor to viscoelastic behaviour [14]. As discussed in the main text, network descriptions that neglect excluded volume appear to be more appropriate for systems closer to the isostatic point.

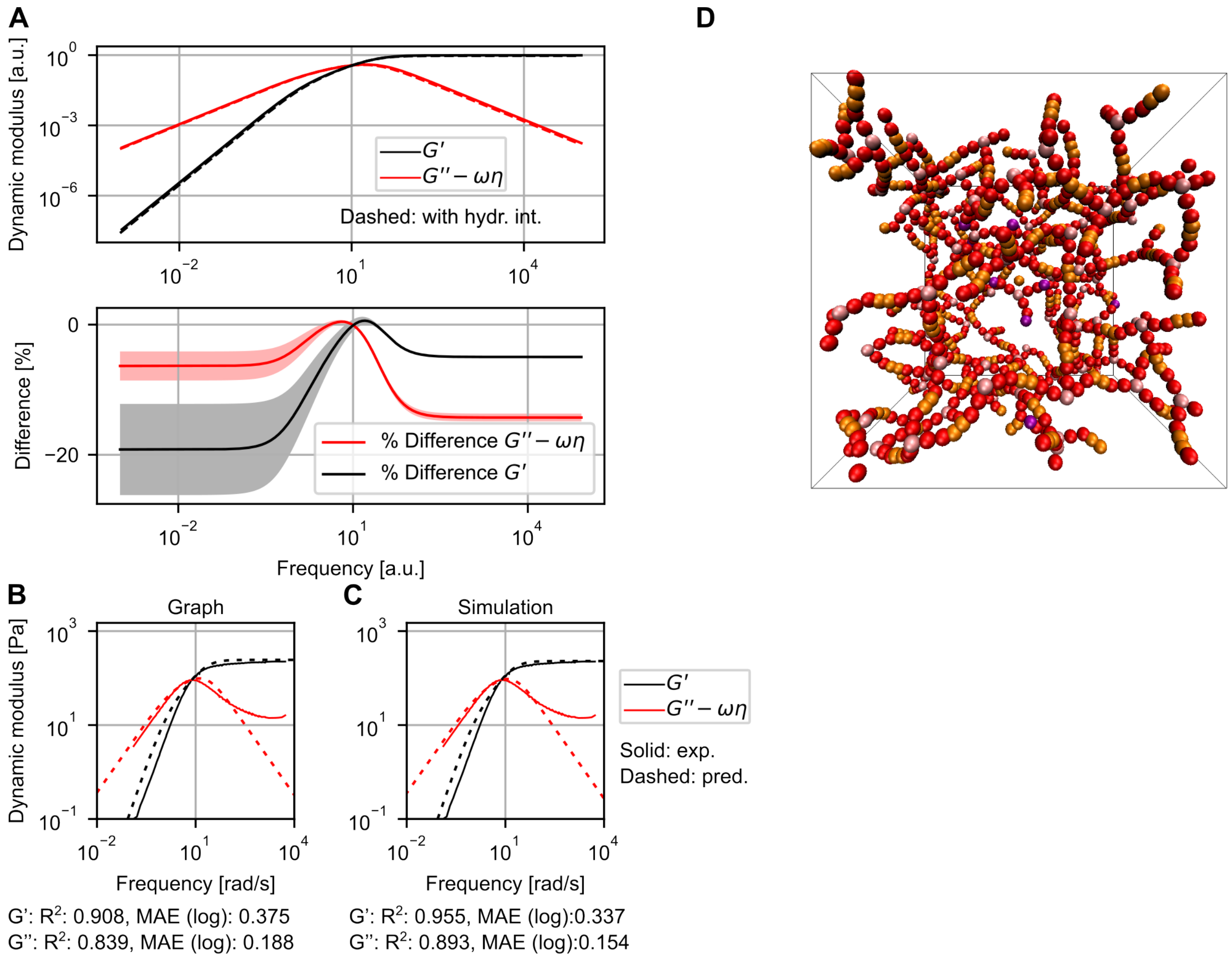


Figure S9: Dynamic moduli predicted by evaluating simulations of 3-armed nanomotifs. (A) Prediction of storage and loss modulus based on the connectivity in simulation of 3-armed nanostars. Dashed lines indicate predictions that account for hydrodynamic interactions, assuming an effective radius of 3.525 nm. Difference in percent shown underneath. (B) Predictions from theoretical graph compared to experimental data. [10] (C) Predictions from simulations accounting for hydrodynamic interactions. $R^2$ and mean absolute error in log space (MAE) shown underneath. MAE values for G" where only integrated up to 50 rad/s, excluding the region with systematic deviations. (D) Snapshot of a simulation with 3-armed DNA nanostars.

#### S7 Availability of code and data

The code developed for this work is maintained and publicly available on GitHub (`https://github.com/aaron-gad/DNA_Nanomotif_ML_MD`). All notebooks and scripts used for this work were also deposited on zenodo (`https://doi.org/10.5281/zenodo.20022811`) and are publicly available. Trajectories of the oxDNA reference simulations for 4-armed nanostars, ReaDDy parameter combinations used for validation, optimisation simulations in ReaDDy as well as all trajectories evaluated for networking and rheological properties in the main text were uploaded to zenodo (`https://doi.org/10.5281/zenodo.20022811`) and are publicly available.

#### References

1 Sengar, A., Ouldridge, T. E., Henrich, O., Rovigatti, L. & Šulc, P. A primer on the oxDNA model of DNA: when to use it, how to simulate it and how to interpret the results. *Frontiers in Molecular Biosciences* **8**, 693710 (2021).

2 Poppleton, E. *et al.* oxDNA: coarse-grained simulations of nucleic acids made simple. *Journal of Open Source Software* **8**, 4693 (2023).

3 Kaufhold, W. oxDNA tutorial (2020). URL `https://github.com/WillTKaufhold1/oxdna-tutorial`.

4 Lukacs, G. L. *et al.* Size-dependent DNA mobility in cytoplasm and nucleus. *Journal of biological chemistry* **275**, 1625–1629 (2000).

5 Kestin, J., Sokolov, M. & Wakeham, W. A. Viscosity of liquid water in the range- 8 c to 150 c. *Journal of physical and chemical reference data* **7**, 941–948 (1978).

6 Einstein, A. Über die von der molekularkinetischen theorie der wärme geforderte bewegung von in ruhenden flüssigkeiten suspendierten teilchen. *Annalen der physik* **4** (1905).

7 Von Smoluchowski, M. Zur kinetischen theorie der brownschen molekularbewegung und der suspensionen. *Annalen der physik* **326**, 756–780 (1906).

8 Snodin, B. E. *et al.* Introducing improved structural properties and salt dependence into a coarse-grained model of DNA. *The Journal of Chemical Physics* **142**, 234901 (2015).

9 Wu, Y.-Y., Bao, L., Zhang, X. & Tan, Z.-J. Flexibility of short DNA helices with finite-length effect: from base pairs to tens of base pairs. *The Journal of Chemical Physics* **142**, 125103 (2015).

10 Conrad, N., Kennedy, T., Fygenson, D. K. & Saleh, O. A. Increasing valence pushes DNA nanostar networks to the isostatic point. *Proceedings of the National Academy of Sciences* **116**, 7238–7243 (2019).

11 Hoffmann, M., Fröhner, C. & Noé, F. ReaDDy 2: Fast and flexible software framework for interacting-particle reaction dynamics. *PLoS Computational Biology* **15**, e1006830 (2019).

12 Sato, Y., Sakamoto, T. & Takinoue, M. Sequence-based engineering of dynamic functions of micrometer-sized DNA droplets. *Science Advances* **6**, eaba3471 (2020).

13 Cohen, S. R., Banerjee, P. R. & Pappu, R. V. Direct computations of viscoelastic moduli of biomolecular condensates. *The Journal of Chemical Physics* **161**, 0951030 (2024).

14 Palombo, G., Weir, S., Michieletto, D. & Gutiérrez Fosado, Y. A. Topological linking determines elasticity in limited valence networks. *Nature Materials* **24**, 454–461 (2025).

15 Rouse Jr, P. E. A theory of the linear viscoelastic properties of dilute solutions of coiling polymers. *The Journal of Chemical Physics* **21**, 1272–1280 (1953).

16 Zimm, B. H. Dynamics of polymer molecules in dilute solution: viscoelasticity, flow birefringence and dielectric loss. *The Journal of Chemical Physics* **24**, 269–278 (1956).

17 Doi, M., Edwards, S. F. & Edwards, S. F. *The theory of polymer dynamics*, vol. 73 (Oxford University Press, 1988).

18 Biffi, S. *et al.* Phase behavior and critical activated dynamics of limited-valence DNA nanostars. *Proceedings of the National Academy of Sciences* **110**, 15633–15637 (2013).
